# Targeting RET in Brain Metastases from Estrogen Receptor Positive Breast Cancer

**DOI:** 10.1101/2025.04.04.647085

**Authors:** Simeng Liu, Geoffrey Pecar, Ye Cao, Fangyuan Chen, Abdalla Wedn, Jennifer M. Atkinson, Jagmohan Hooda, Steffi Oesterreich, Adrian V. Lee

## Abstract

Brain metastases (BrM) occur in 10-15% of patients with estrogen receptor (ER)-positive breast cancer, and remain a significant clinical challenge. While current therapeutic paradigms, including surgical resection and systemic therapy, have efficacy in the management of BrM from ER+BC, median overall survival following the diagnosis of BrM is approximately 18 months. The limited efficacy of current therapies, along with a relative paucity of therapeutic options, highlight the urgent clinical need to identify druggable targets for BrM from ER+BC.

We previously identified recurrent overexpression of the receptor tyrosine kinase RET in breast cancer BrM relative to patient-matched primary tumors. The principal ligand for RET, Glial Cell-Derived Neurotrophic Factor (GDNF), is predominately expressed in the brain, where it functions in the maintenance of neurons and multiple glial cell types. Here, we show that increased RET expression and activation correlates with brain metastasis in breast cancer patients. Further, we confirm the role of RET signaling in the promotion of metastatic properties *in vitro,* including tumor spheroid formation, migration, invasion, and transformation. Using RET-overexpressing and RET-deficient models in four ER+BC cell lines, we demonstrate that RET overexpression enhances activation of RET downstream targets, such as ERK and AKT, and promotes metastatic phenotypes in *in vitro*, *ex vivo* and *in vivo* assays. Importantly, we demonstrate that the phenotypic effects of GDNF-mediated RET signaling can be abrogated by administration of Pralsetinib, a highly potent RET-selective kinase inhibitor. Using an *ex vivo* mouse brain slice co-culture model, we further demonstrate that RET overexpression in breast cancer cell lines potentiates their colonization and invasion into the brain tissues, which are reduced in RET-knockdown cell lines. RET overexpression facilitates brain colonization of breast cancer cells *in vivo* and is associated with reduced survival. Using proteomic analyses, we identify novel GDNF-mediated signaling pathways in breast cancer cell line models, including PLCγ, P70S6K, STAT3 and CREB. RET-mediated activation of this range of downstream targets is sensitive to inhibition by Pralsetinib. In sum, this study identifies and characterizes RET as a targetable driver of breast cancer brain metastasis, and further describes the action of Pralsetinib in ER-positive breast cancer cell lines, providing insights to improve current targeted therapies for patients with breast cancer brain metastasis.

**Bullet points:**

- RET expression and activation are increased in breast cancer brain metastases.
- Pralsetinib potently inhibits RET activation in RET overexpressing and in non-overexpressing breast cancer cell line models.
- RET overexpression and activation confers pro-metastatic properties including cell proliferation, colonization, migration and invasion in vitro and in *ex vivo* brain co-culture and enhances brain colonization of breast cancer cells *in vivo*.
- GDNF increases activation of ERK1/2, AKT, PCLγ, P70S6K, STAT3, and CREB in RET overexpressing MCF-7 cells.
- CREB or P70S6K knockdown attenuates GDNF stimulated activation of RET downstream targets and cell migration.

**Graphical Abstract:** 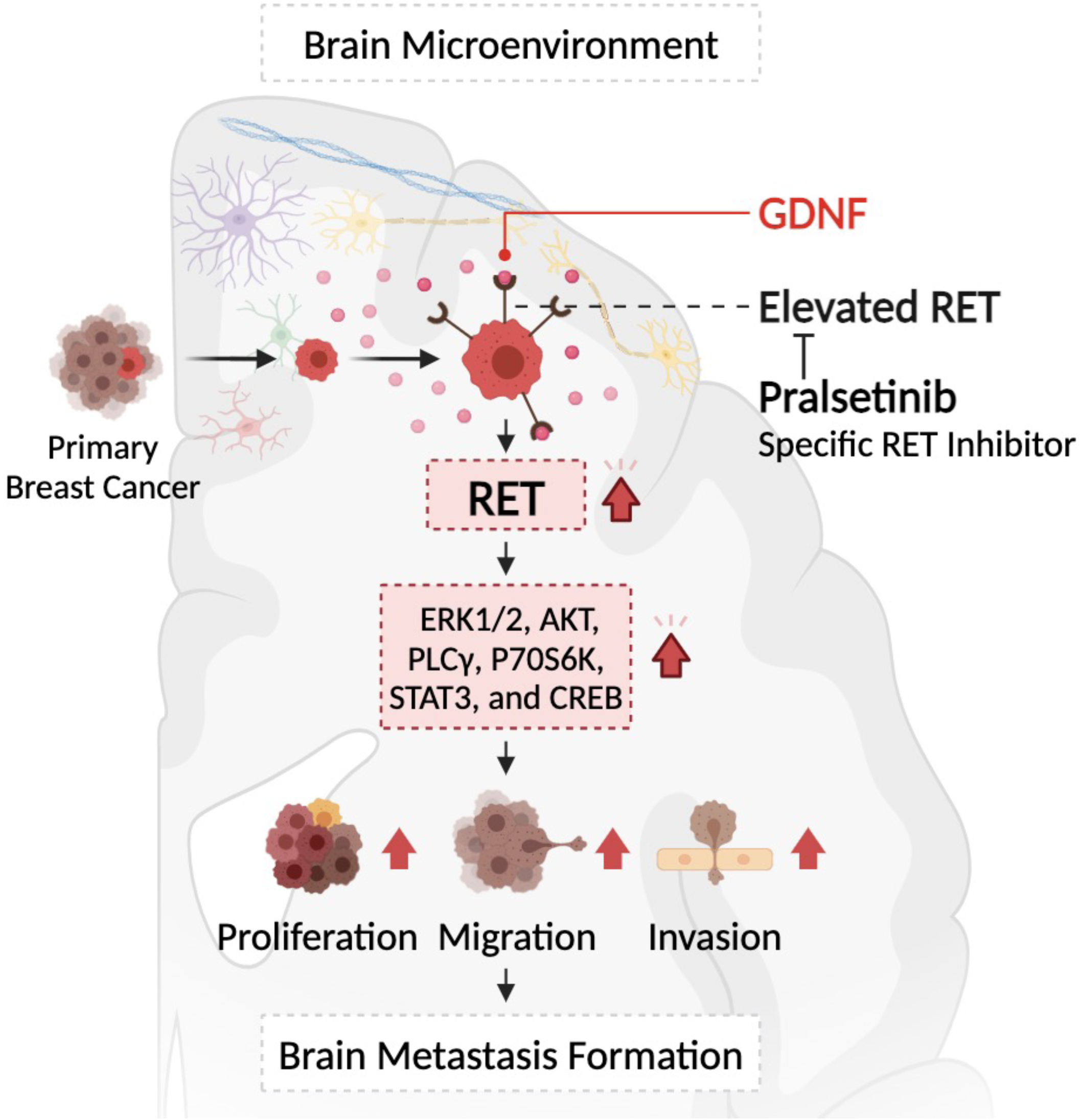

## Introduction

Brain metastases (BrM) affect approximately 10-15% of patients with estrogen receptor (ER)-positive breast cancer (ER+BC), and remain a major clinical challenge [1]. Current therapeutic strategies consisting primarily of surgical resection, cytotoxic chemotherapy, and radiation have efficacy in the management of BrM. However, median overall survival following the diagnosis of BrM from ER+BC remains approximately 18 months [2], making the identification of additional therapeutic targets an unmet clinical need. While multiple systemic therapies are under active investigation for the management of BrM from ER+BC, strategies of interest are dominated by ER and cyclin-dependent kinase inhibition [3]. Using genome-wide exome capture RNA sequencing, we have previously identified recurrent overexpression of the receptor tyrosine kinase (RTK) RET in breast cancer BrM relative to patient-matched primary tumors [4]. The principal ligand for RET, Glial Cell-Derived Neurotrophic Factor (GDNF), is highly expressed in the brain and peripheral nervous tissue, where it functions in the maintenance of both neurons and glial cells [5]. Recently, RET has been shown to drive adhesion and *ex vivo* brain colonization in organoid models, providing further evidence for a role in the establishment of secondary CNS lesions [6]. Here, we expand on the current understanding of a role for RET in BrM from ER+BC by describing novel signaling interactions and demonstrating the effects of RET overexpression and knockdown in *in vivo* BrM growth.

RET is a single-pass transmembrane RTK that functions primarily in the development, maturation and maintenance of the kidney and the enteric nervous system, and has recently been shown to regulate phenotypic changes associated with lactation and mammary duct involution [7, 8]. Activation of RET signaling requires binding of GDNF family ligands (GFLs) to GDNF family receptor-α (GFRα) family co-receptors. The subsequent formation of ternary GFL-GFR-RET complexes triggers RET dimerization and kinase domain phosphorylation, which in turn induces the phosphorylation of numerous downstream targets including ERK, AKT, and PLCγ-1 [7]. The *RET* proto-oncogene contributes to tumor formation through two major mechanisms, gain-of-function mutations (which include both point mutations and gene fusions that lead to constitutive kinase activity) and increased expression of wild-type receptors [7]. Aberrant *RET* expression is associated with several tumor types including medullary thyroid carcinoma (MTC), papillary thyroid carcinoma (PTC), non-small cell lung carcinoma (NSCLC), and breast cancer [9–11]. RET and GFRα1 are expressed in approximately 30–70% of human breast cancers [12]. Among breast cancer subtypes, RET expression is most common in ER+ tumors, which comprise approximately 70% of all breast tumors [13–15]. In patients with ER+ breast cancer, RET expression is elevated in recurrent tumors compared to normal tissues or corresponding primary lesions, and is correlated with larger tumor size, more advanced tumor stage, and reduced metastasis-free and overall survival [13, 14, 16, 17]. Previous studies have demonstrated that activation of RET signaling promotes breast cancer cell proliferation, survival, and migration *in vitro* [14, 16, 18]. While the function of RET in the formation of primary breast tumors and the development of resistance to endocrine therapy has been extensively studied, the role of aberrant RET expression and activation in breast cancer BrM is less clearly understood.

RET is a target of multiple multi-kinase inhibitors (MKIs), including Cabozantinib and Vandetanib, which inhibit RET less potently than their principal target kinases such as VEGFR1/2/3, MET, or EGFR [19, 20]. Increased recognition of the function and prevalence of *RET* alterations in multiple cancers has led to the development of the first generation of RET-selective kinase inhibitors (Selpercatinib and Pralsetinib), which are currently clinical investigation for multiple cancer types and have shown potent preclinical activity against RET mutations [21–23]. The potency of RET blockade by Pralsetinib has been demonstrated in NSCLC and MTC cell lines with constitutively active RET mutations, showing 10 - 100-fold reductions in IC_50_ relative to MKIs [21], as well as clinical activity in lung and thyroid carcinoma patients (NCT03037385). Notably, Pralsetinib and Selpercatinib have been shown to generate a therapeutic benefit in intracranially implanted NSCLC cell lines and in BrM in patients harboring *RET* fusions [21, 24, 25]. This suggests that the current generation of RET-selective inhibitors may accumulate beyond the blood-brain barrier at therapeutic concentrations, making them a valuable potential strategy for the management of BrM.

In this study, we provide evidence that increased activation of RET correlates with the incidence of brain metastasis in breast cancer patients. We demonstrate that RET overexpression amplifies activation of RET downstream targets, including ERK and AKT, and promotes multiple pro-metastatic properties. We demonstrate that the RET-selective kinase inhibitor Pralsetinib is highly effective and potent in the abrogation of GDNF-mediated RET signaling and in the reduction of pro-metastatic phenotypes. *Ex vivo* brain slice co-culture models reveal that RET overexpression in breast cancer cell lines potentiates colonization and invasion in the brain microenvironment. Furthermore, *in vivo* data show that RET overexpression facilitates brain colonization of breast cancer cells and is associated with poor overall survival of the xenografts. In summary, this study identifies and characterizes RET as a targetable driver of breast cancer brain metastasis, and further describes the action of Pralsetinib in ER-positive breast cancer cell lines, providing insights to improve current targeted therapies for patients with breast cancer brain metastasis.

## Results

### RET expression and activation is increased in breast cancer brain metastases

Activation of RET by GDNF-GFRα complexes initiates multiple downstream pathways involved in promotion of cell proliferation and survival to facilitate tumor initiation and progression [7]. Given that the levels of RET expression are elevated in breast cancer BrM, we sought to further analyze the relationship between the activation of RET-mediated downstream signaling and brain metastasis. To this end, we utilized a proliferation-independent GDNF-responsive gene set (GDNF-RGS) developed by Morandi et al. [26] to examine RET expression and activation of the GDNF-RET signaling axis in RNAseq datasets of both primary and metastatic ER+ breast tumors.

Considering the extensive functional interactions between the RET and ER pathways [15, 16], we analyzed our previously reported cohort consisting of 67 ER-positive (ER+) clinical samples that contains 35 metastatic tumors from four metastatic sites (brain, bone, gastrointestinal tract, and ovary) and the corresponding primary tumors. RET expression is higher in metastatic brain tumors than either bone or ovarian metastases, with the highest median (Fig. 1a) among all four metastatic sites. Although no statistical significance in GSVA scores of GDNF-RGS was observed between tumors from different metastatic sites, BrM lesions exhibited the highest median level of GDNF-RGS (Fig. 1b). Within the 9 pairs of ER+ primary tumor patient-matched BrM, RET expression was increased in 7 cases, with a concurrent increase in GDNF-RGS expression in 5 cases (Fig. 1c), demonstrating a recurrent gain in both expression and activation of RET in BrM lesions. Consistently, analyses of multiplatform genomic profiles of matched brain-metastatic and primary breast tumors from AURORA [27] revealed the highest median of RET expression and GSVA score in brain metastatic tumors among all metastatic sites (Fig. S1a-b). Higher median RET expression was observed in primary tumors corresponding to BrM lesions than other metastatic sites (Fig. S1c-d).

**Figure 1.**
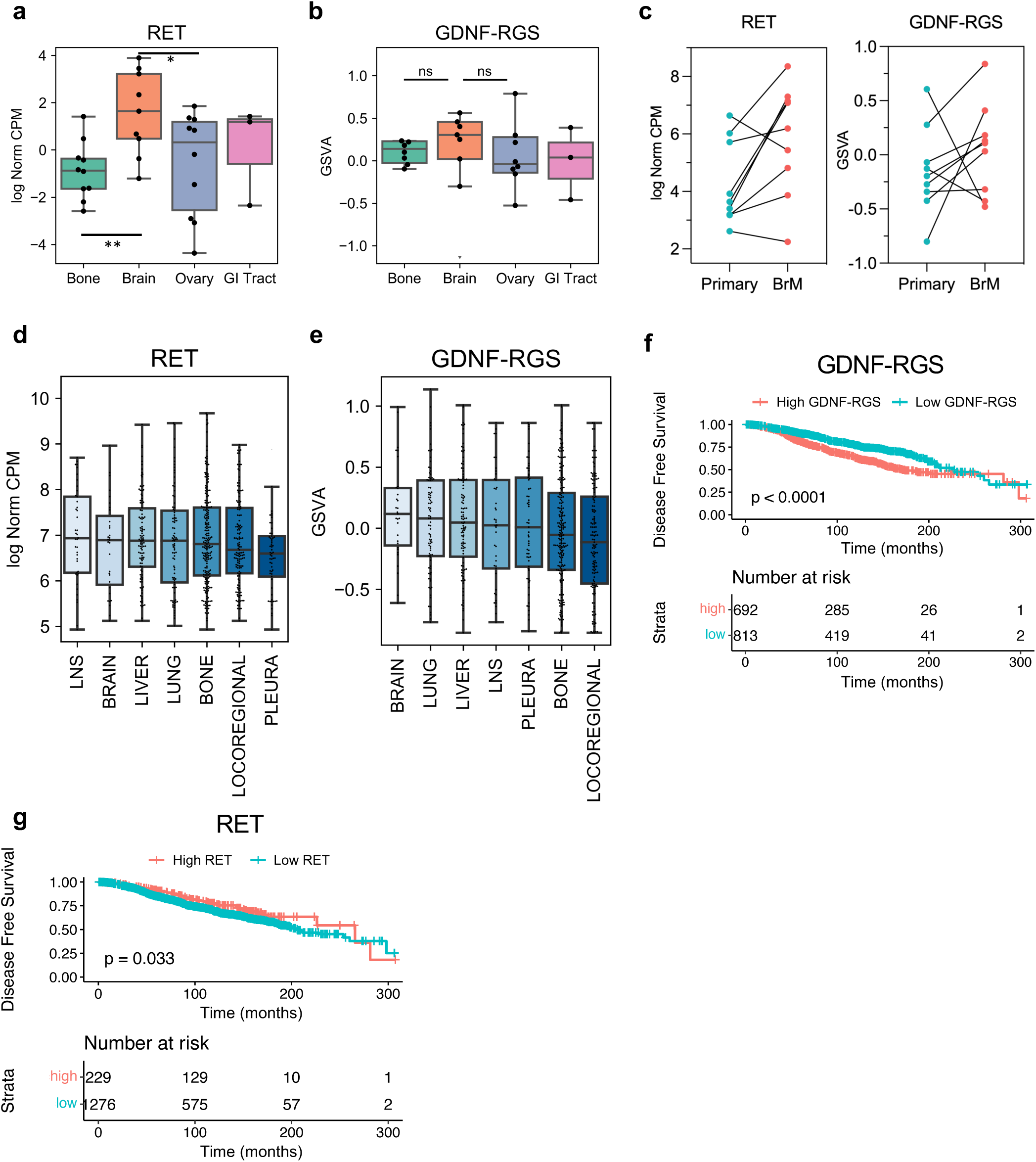
RET expression and signaling was increased in breast cancer brain metastasis. (a) RET expression (log2NormCPM:log2 normalized counts per million) in ER+ breast cancer among four metastatic sites of in-house dataset. Wilcoxon test. (b) GSVA score of GDNF-RGS signature in ER+ breast cancer among four metastatic sites of in-house dataset. (c) Paired ladder plot of RET expression (left) and GDNF/RGS GSVA score (right) in BrM versus matched primary tumor. (d) RET expression (log2NormCPM) in ER+ primary tumors in in-house dataset. (e) GSVA score of GDNF-RGS signature in ER+ primary tumors in METABRIC. Future metastatic sites were shown. (f) Kaplan-Meier (KM) plot of disease free survival (DFS) with high/low GDNF-RGS GSVA scores in ER+ cases of METABRIC. High/low GNDF-RGS score was determined based on an outcome-oriented methods providing a value of a cutpoint that correspond to the most significant relation with survival. p<0.0001, log rank test. (g) KM plot of DFS with high/low RET expression in ER+ cases of METABRIC.

Next, we examined the relationship between RET expression and activation and the clinical outcomes of breast cancer patients. We analyzed the METABRIC dataset with a record of histopathology and RNAseq data of primary tumors grouped by their future metastatic sites and disease-free survival data acquired through follow-up. Consistent with our cohort, BrM lesions showed the highest median GDNF-RGS scores, and the second-highest median levels of RET expression among metastatic sites with 20 or more available samples (Fig. 1d,e). Increased GDNF-RGS was significantly associated with poor survival (Fig. 1f). Interestingly, RET expression was significantly associated with increased disease-free survival (p=0.033) (Fig. 1g). Neither GDNF-RGS nor RET expression were predictive of BrM-free survival within the ER+ cases with subsequent BrM formation (Fig. S1g-h), however, this may be due to reduced cohort size. Taken together, we found a general elevation of RET expression in BrM relative to the matched primary breast tumor and other metastatic sites.

### Pralsetinib potently inhibits RET activation in RET-overexpressing and in non-overexpressing breast cancer cell line models

Pralsetinib, a RET-selective kinase inhibitor, has shown highly effective RET blockade in cell line models of NSCLC and MTC [21]. In contrast, the efficacy of Pralsetinib in breast cancer models remains largely unexamined. To explore the relevance of the GDNF-RET signaling axis in breast cancer, a cohort of 60 breast cancer cell lines was screened for mRNA levels of RET, the coreceptor GFRα1 and the ligand GDNF, from which 4 representative models were selected for further study. The cell lines selected comprise 2 models of invasive breast carcinoma (no special type), and 2 models of invasive lobular carcinoma (ILC) (Fig. 2a), representing RET-positive models of the two most common breast cancer histologic subtypes. The T-47D cell line displayed undetectable levels of RET (Fig. 2b,f) and GFRα1 [14, 16], and therefore served as a negative control. GDNF expression was found to be minimal across all 4 cell lines, suggesting no role for endogenous ligand expression. We next assessed the effect of exogenous GDNF on each cell line model. After overnight serum starvation in low (0.5%) serum medium followed by 30 min treatment with 100 ng/mL GDNF, RET phosphorylation at Y905 was significantly upregulated in MCF-7 and SUM44 cell lines, but was minimally upregulated in the MM134 cell line (Fig. 2b). T-47D cells displayed no significant change in RET, ERK, or AKT phosphorylation in response to GDNF treatment, presumably due to the absence of GFRα1 co-receptor expression and low expression of RET. shRNA-mediated RET knockdown in the MCF-7 and SUM44 cell lines significantly reduced both RET expression and phosphorylation in response to GDNF with a concurrent reduction in downstream ERK and AKT phosphorylation, indicating specificity of GDNF-induced RET signaling (Fig. 2c-d). However, RET knockdown was incomplete in the SUM44 cell line, in which GDNF-mediated phosphorylation of ERK and AKT persisted in the RET-deficient model (Fig. 2d). Pretreatment with Pralsetinib exhibited stronger abrogation of GDNF-RET mediated signaling than RET knockdown in both MCF-7 and SUM44 cell lines (Fig. 2c-d). To examine the effects of RET signaling in the T-47D and MM134 cell lines, which exhibit minimal responses to GDNF treatment, we expressed exogenous RET in these two cell lines. In both models, phosphorylation of RET and multiple downstream targets in response to GDNF was significantly increased after RET overexpression, and was potently inhibited by Pralsetinib (Fig. 2e-f). RET (Y905) and ERK exhibited ligand-independent phosphorylation in MM134 RET overexpressing (RET-OE) cells, consistent with reports of ligand-independent RET and MEK activation in NIH-3T3 cells following exogenous RET expression due to autophosphorylation of the receptor [11, 12]. Additionally, strong autophosphorylation of RET was observed in MM134 RET-OE cells, which persisted at Y905 in the presence of Pralsetinib. Treatment with recombinant GFRα1 facilitated GDNF-induced RET signaling in T-47D cells (Fig. 2f), consistent with the restricted co-receptor expression in this specific cell line. Interestingly, T-47D RET-OE cells were more sensitive to Pralsetinib-mediated abrogation of ERK1/2 and AKT phosphorylation than control cells transfected with an empty vector (EV). MCF-7 was selected as the most appropriate cell line model for subsequent *in vitro* studies.

**Figure 2.**
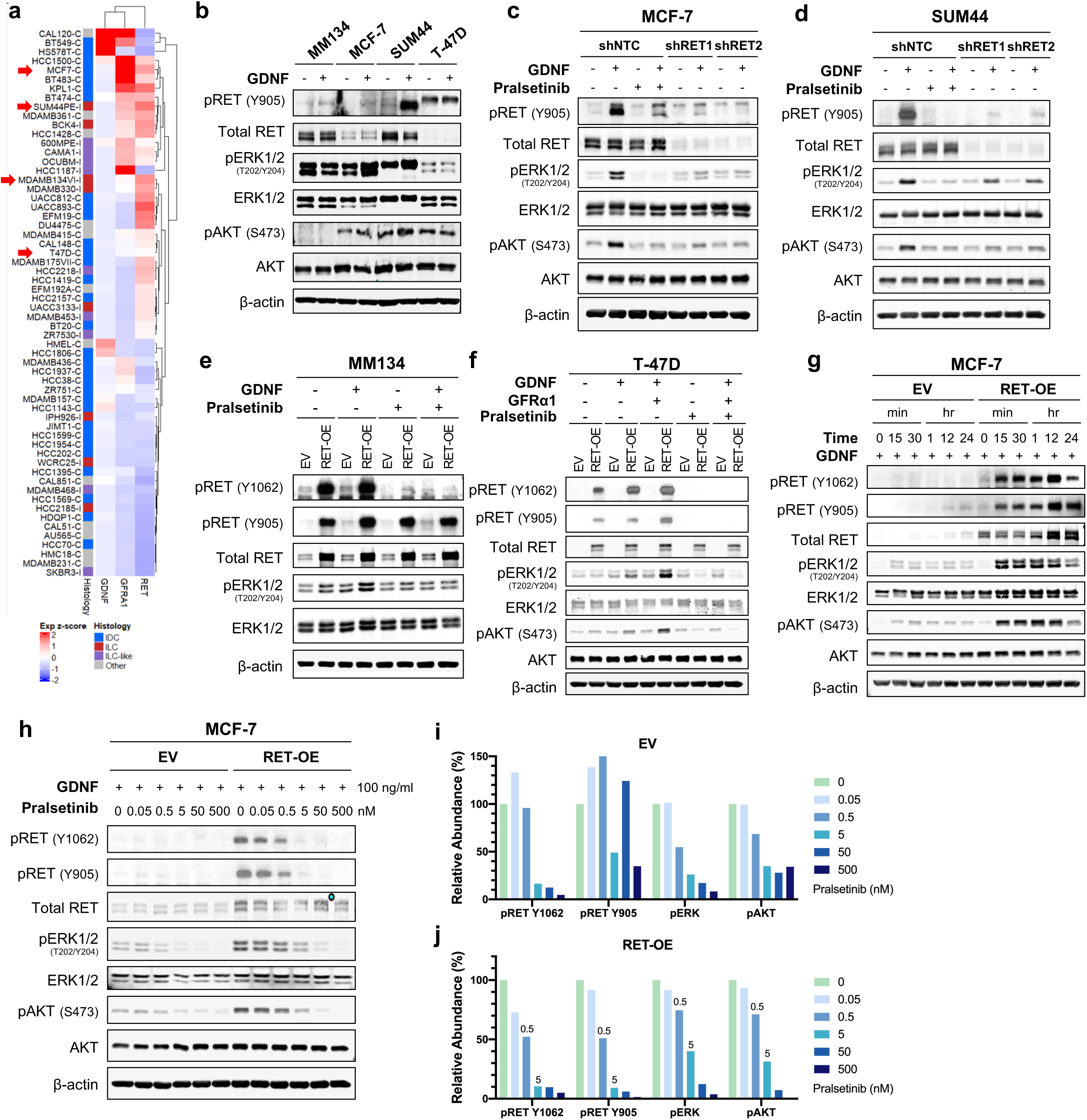
Pralsetinib potently inhibits RET activation in breast cancer cell line models. (a) mRNA levels of RET, GFRα1 and GDNF level in CCLE (C) and ICLE (I). Arrows indicate cell line models involved in this study. (b-g). Immunoblotting of the indicated breast cancer cell line(s). (h-j) Immunoblotting (h) and quantification (i-j) of pralsetinib dose response on RET and downstream targets in MCF-7 cells. Relative abundance was calculated by normalization to intensity of β-actin measured in Image J. GDNF, 100 ng/ml, 30 min. GFRα1, 10 ng/ml, 30 min. Pralsetinib, 500 nM, 16 hr. Vehicle, distilled water for GDNF, PBS for GFRα1, and DMSO for Pralsetinib. EV; empty vector transfected cells. RET-OE; RET overexpressing cells. shNTC; cells transfected with non-targeting control shRNA. shRET; cells transfected with shRNA targeting hRET.

The relationship between duration of GDNF treatment and activity of RET signaling was examined in MCF-7 cells with basal (EV) or elevated (RET-OE) RET expression (Fig. 2g). RET phosphorylation showed a rapid response to GDNF in MCF-7 RET-overexpressing cells, with phosphorylation of RET and downstream ERK and AKT significantly elevated after 15 minutes of GDNF treatment. RET activation peaked after 12 hours of treatment, and persisted through 24 hours in the MCF-7 RET-OE model. A similar pattern was observed in the SUM44 model, though peak RET activity was achieved at 30 minutes of treatment, followed by a sudden drop after 1 hour (Fig. S2a). These results indicate that GDNF has a rapid and durable effect on MCF-7 cells, which serve as the representative RET+ breast cancer cell line in the remainder of these studies.

Dose-response assays were used to examine the efficacy and potency of Pralsetinib-mediated inhibition of GDNF-RET signaling. Pralsetinib has demonstrated ≥10-fold increased potency over approved MKIs against oncogenic RET variants, with over 50% inhibition of RET and AKT activation at concentrations as low as 5 nM [21]. Here, we assessed the activity of Pralsetinib against RET signaling in breast cancer models with elevated RET expression. We found that the RET-selective inhibitor reduced GDNF-mediated phosphorylation of RET at both Y1062 and Y905 with an IC_50_ of 0.5 nM after 16 hr treatment, reaching a 90% of total inhibition efficacy at a concentration of 5 nM (Fig. 2h-j). Pralsetinib also decreased phosphorylation of ERK1/2 and AKT at 5 nM, though phosphorylation of ERK1/2 was less sensitive to inhibitor treatment in RET-OE cells relative to empty vector control (Fig. 2h). Activation of both ERK and AKT was completely blocked by treatment of Pralsetinib at a concentration of 500 nM (Fig. 2j). The SUM44 cell line was examined, as a GDNF-responsive ILC model. Pralsetinib reduced phosphorylation of RET at 0.5 nM in SUM44 cells (Fig. S2b). Similar to findings in MCF-7, in SUM44 Pralsetinib completely suppressed activation of ERK1/2 at a concentration of 500 nM (Fig. S2c). Interestingly, phosphorylation of AKT in SUM44 cells persisted at an intermediate level despite increasing Pralsetinib concentration, suggesting an alternative pathway activating AKT in this specific cell line. Collectively, these results demonstrate that Pralsetinib effectively abrogates GDNF-mediated activation of RET and downstream signaling pathways in RET-overexpressing breast cancer cell line models.

### RET overexpression and activation confer pro-metastatic properties *in vitro*

GDNF-RET signaling has previously been shown to contribute to metastatic phenotypes through the facilitation of cell proliferation, colony formation, migration, and invasion. GDNF treatment significantly enhanced 2D cell proliferation in both MCF-7 EV and RET-OE cell lines (Fig. 3a). However, RET overexpression did not result in increased GDNF-dependent proliferation, suggesting that basal RET expression in MCF-7 cells is sufficient to drive GDNF-dependent proliferative phenotypes [29, 30]. Treatment with Pralsetinib at 500 nM completely abrogated the stimulatory effect of GDNF on 2D proliferation in both EV and RET-OE cell lines (Fig. 3a). Cells with shRNA-mediated RET-knockdown showed no response to GDNF in 2D culture, however, transfection with non-targeting control shRNA (shNTC) also eliminated the response to GDNF, indicating a RET-unrelated effect of shRNA transfection (Fig. S3a). The effect of GDNF-RET signaling on cell proliferation was also evaluated in Ultra-Low Attachment (ULA) culture, which minimizes the effects of cell attachment and contact by facilitating suspended growth over a hydrogel surface [31]. Consistent with the observation in 2D culture, GDNF treatment increased cell proliferation in ULA culture in both EV and RET-OE cell lines, although RET overexpression did not enhance the stimulatory effects of GDNF on the proliferative phenotype (Fig. 3b). GDNF-stimulated cell proliferation was decreased by shRNA-mediated RET knockdown, confirming that GDNF promotes cell growth of MCF-7 cells through a RET-dependent mechanism (Fig. S3b). Pralsetinib effectively abrogated GDNF-promoted cell growth in all groups. Taken together, these data demonstrate that GDNF promotes cell proliferation both in 2D culture and in anchorage-independent conditions. Pralsetinib treatment abrogated the stimulatory effect of GDNF treatment in breast cancer models with either basal or overexpressed RET.

**Figure 3.**
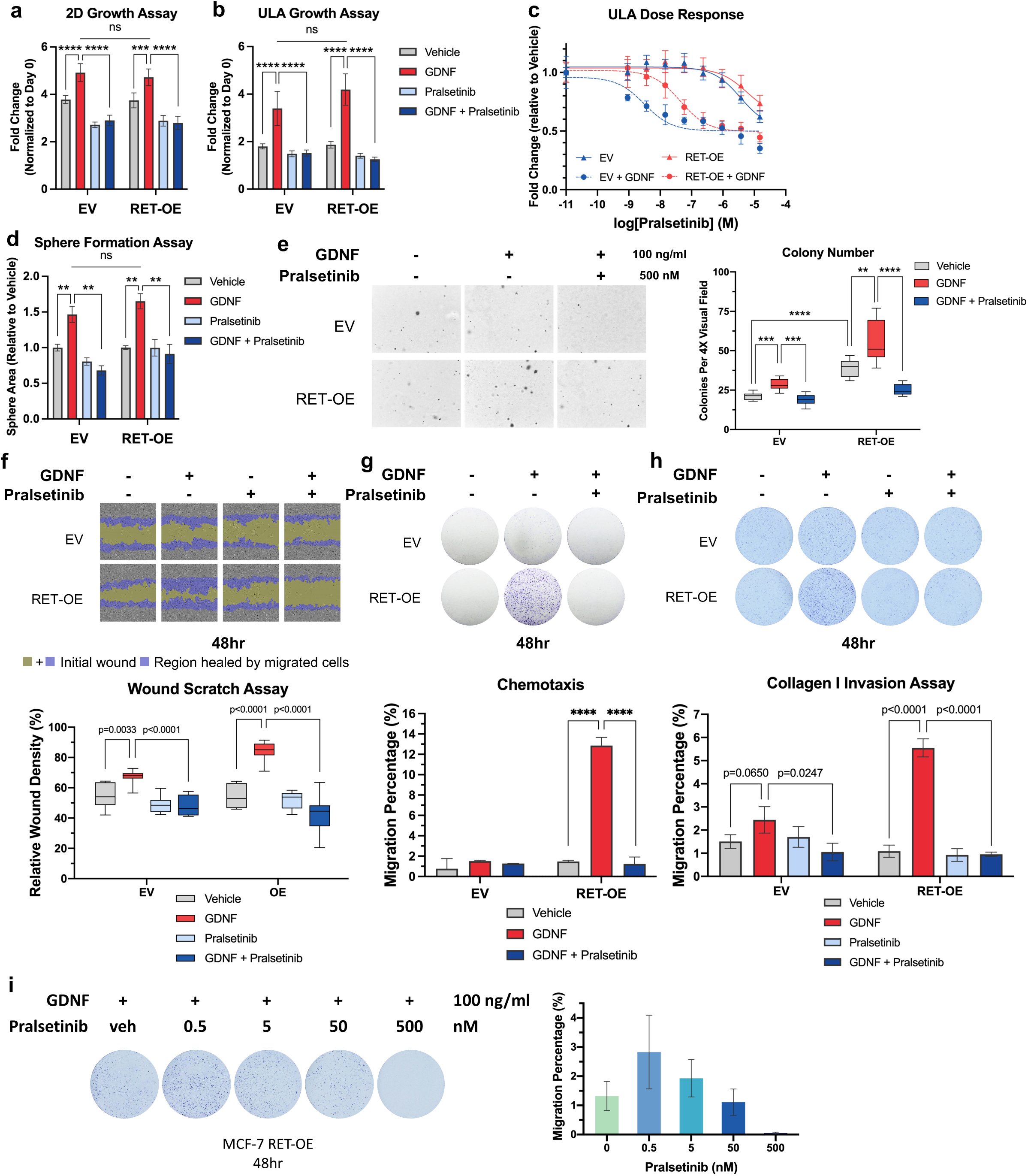
RET overexpression and activation confer pro-metastatic properties *in vitro* in MCF-7 RET-overexpressing model. (a-b) Cell growth assay in 2D (a) and ULA (b) conditions in MCF-7 RET overexpressing models. (c) Cell survival assay to determine dose response of Pralsetinib in MCF-7 cells in ULA culture. (d) Quantification of sphere formation assay. (e) Representative images and quantification of soft agar assay. GDNF promoted cell growth in RET overexpressing MCF-7 cells, which was inhibited by treatment of Pralsetinib. (f) Wound scratch assay. Mitomycin C (5 ug/ml) was used to prohibit cell proliferation. Top, 10X images are taken in Incucyte with initial wound masked with yellow and healed region masked with purple. Bottom, relative wound density was quantified using Incucyte wound scratch module, reflecting portion of region healed by migrated cells to initial wound. Results at 48 hr were plotted. (g-h) Images and quantification of crystal violet–stained transwell inserts in (g) chemotaxis and (h) rat tail collogen I invasion assays. Multiple t test was performed (FDR = 0.05). (i) Boyden transwell assay to determine dose response of Pralsetinib on chemotaxis to GDNF in MCF-7 RET-OE cell. Representative images and quantification was shown. GDNF, 100 ng/ml. Pralsetinib, 500 nM. Vehicle, distilled water for GDNF, and DMSO for Pralsetinib. EV; empty vector transfected cells. RET-OE; RET overexpressing cells. *, 0.01 < p < 0.05, **, 0.001 < p < 0.01, ***, 0.0001< p < 0.001, ****, p < 0.0001.

The potency of Pralsetinib in the inhibition of ULA cell proliferation was further assessed in an effort to characterize the inhibitor’s properties in breast cancer cell line models. In the absence of GDNF, no significant difference in the response to Pralsetinib treatment was observed between MCF-7 EV and RET-OE cells, with IC_50_ values of approximately 5 μM calculated for both cell lines (Fig. 3c, Fig. S2d), a relatively high concentration to suppress cell growth. In the presence of GDNF, IC_50_ was markedly reduced in both cell lines, with RET-OE cells requiring a 10-fold higher dose for growth suppression (43.4 nM versus 3.12 nM). Given that GDNF is relatively abundant in the brain microenvironment, these data suggest that Pralsetinib may be highly efficient in inhibiting the proliferation of RET-expressing breast cancer cells at low doses.

Formation of colonies improves tumor cell survival away from the primary lesion, thereby facilitating distant metastasis. We next sought to evaluate colony formation using sphere formation assays, which encourages the formation of intercellular contacts via adherens junctions in a ULA environment [32, 33]. GDNF induced the formation of larger spheres in MCF-7 cells with or without exogenous RET expression (Fig. 3d). While RET overexpression did not further promote sphere formation, RET-deficient cells lost response to GDNF stimulation, indicating the specificity of GDNF-RET mediated signaling in promotion of breast cancer cell proliferation and colony formation (Fig. S3c). The stimulatory effect of GDNF was eliminated with both RET knockdown and Pralsetinib treatment. We employed soft agar gel assays to further examine anchorage-independent colony formation [34, 35]. Notably, RET overexpression increased both colony number and colony size in this assay (Fig. 3e, Fig. S3d). GDNF-promoted colony formation was effectively abrogated by Pralsetinib in both cell lines. Importantly, RET-OE cells formed more colonies in soft agar than EV controls in the absence of GDNF, suggesting that RET may function independently of GDNF to promote survival in anchorage-independent conditions.

GDNF has been reported to increase cell motility in various cancers, including breast cancer and glioma [14, 36]. To investigate the role of RET overexpression on cell migration, we utilized wound healing assays, which introduce an artificial gap in a cell monolayer, with migration represented by the rate and extent of wound healing. GDNF significantly promoted cell migration in RET-overexpressing cells compared to the empty vector control, with migration inhibited by Pralsetinib in both cell lines (Fig. 3f). We next utilized Boyden chamber transwell assays as an alternative means to study cell migration, in which GDNF was employed as a chemoattractant (Fig. 3g). While MCF-7 cells with basal RET expression displayed limited chemotaxis toward GDNF, RET overexpression significantly increased chemotaxis, supporting the hypothesis that increased RET expression may facilitate metastatic colonization of a GDNF-rich environment. Pralsetinib treatment abrogates cell migration in this system, with complete inhibition at 500 nM (Fig. 3f, i), further indicating the potential therapeutic utility of RET inhibition.

Parental MCF-7 cells with basal level expression of RET typically exhibit minimal invasiveness, but administration of GDNF has been shown to enhance cell invasion [14, 37]. To assess the effect of RET overexpression on invasion in our cell line model, we employed an optimized Collagen I coated Boyden chamber assay as previously described [14]. In contrast to our studies of chemotaxis, GDNF induced only minor invasion in MCF-EV cells (Fig. 3h). However, GDNF significantly promoted cell invasion in MCF-7 RET-OE cells relative to empty vector control. Pralsetinib abrogated GDNF-promoted cell invasion in both RET overexpressing cells and empty vector control.

Conclusively, these observations demonstrate the stimulatory effect of GDNF-RET signaling on pro-metastatic phenotypes, including cell proliferation, colonization, migration and invasion. Notably, RET overexpression itself did not metastatic properties in MCF-7 cells, highlighting the necessity of GDNF for sufficient RET activation and resulting phenotypic changes.

### RET overexpression drives colonization and invasion of *ex vivo* brain slice cultures

In order to assess the effects of RET expression on colony formation and invasion of brain tissue, we adapted an organotypic coculture system utilizing slices of mouse brain as previously described [38, 39]. RFP-labeled RET-OE cells or EV control cells were cultured on mouse brain slices, which were maintained in enhanced brain culture media (Fig. 4a). GDNF expression in mouse liver is undetectable [40], and therefore liver slices were employed as negative control (Fig. 4b). RET overexpression increased the colonizing MCF-7 number threefold in brain slice co-culture compared to empty vector control, and approximately sixfold relative to liver co-culture (Fig. 4b). To evaluate the effects of RET inhibition on this phenotype, cells were pretreated with Pralsetinib (500 nM) overnight before addition to co-culture systems. Pralsetinib treatment reduced brain slice colonization by MCF-7 RET-OE cells to levels comparable to EV controls (Fig. 4b). These data demonstrate that RET overexpression enhances tissue colonization by breast cancer cells, and that this effect is enhanced in the brain relative to other frequent sites of breast cancer metastasis. Further, RET inhibition effectively reduces colonization of brain tissues, suggesting a role for RET as a druggable target in the management of BrM.

**Figure 4.**
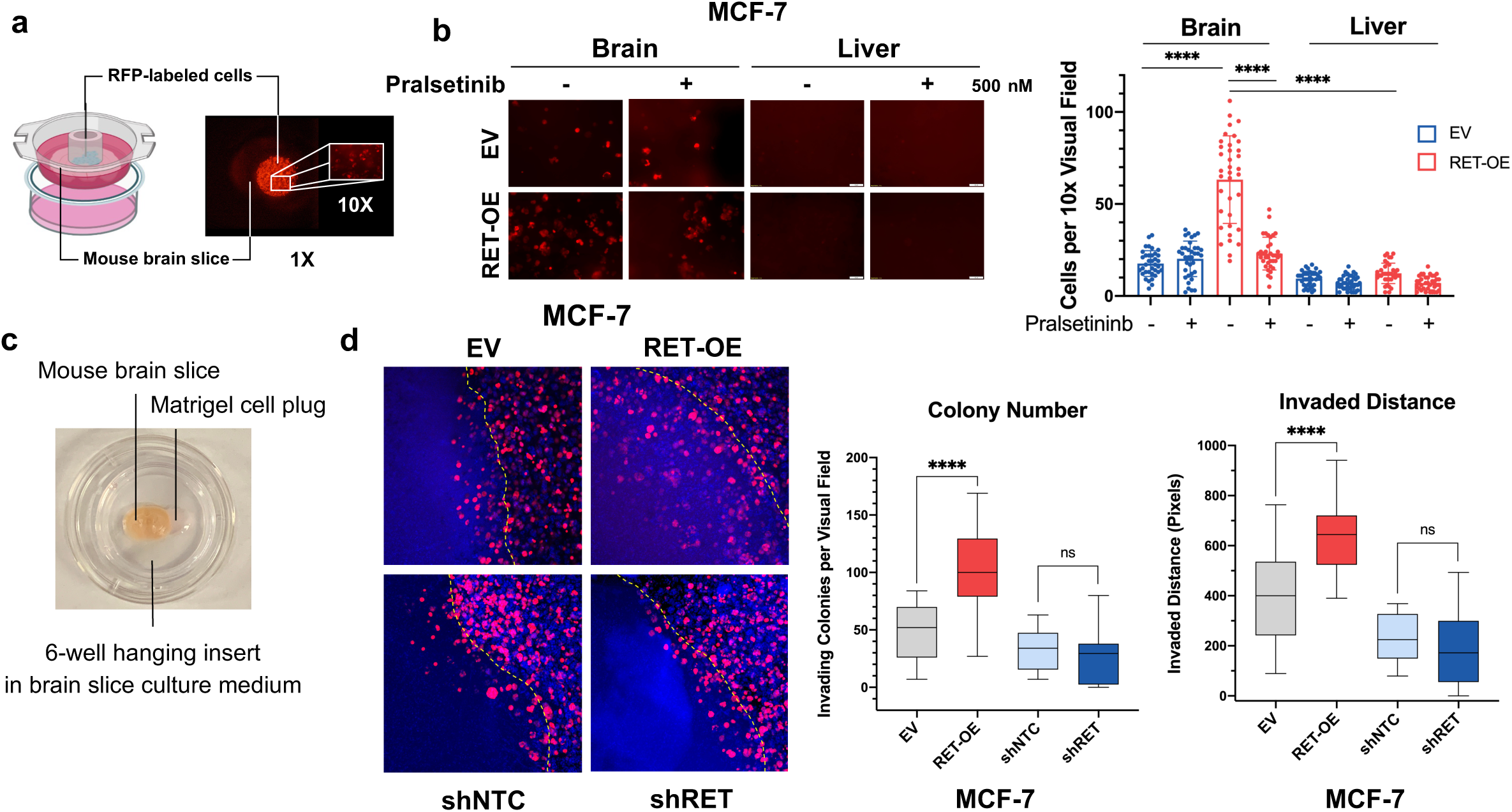
RET overexpression is critical for colonization and invasion of *ex vivo* brain slice cultures. (a) Diagram of *ex vivo* brain slice coculture setup and representative fluorescent image. 50,000 RFP-labelled MCF7 cells were incubated on top of 300-micron brain slices for 5 days before tissues were gently washed in PBS (with calcium and magnesium) for 5 minutes before fluorescent microscope imaging. (b) Left panel: Fluorescent 10X visual fields after 5-minute PBS washing. Right panel: Quantification of cells observed per visual field (10X), ImageJ multipoint tool. Each dataset represents n=36 samples scored between 3 independent experiments. Unpaired t-test. (c) Picture of experiment setup. Mouse brain slice was cultured in 6-well hanging insert on top of brain slice culture medium. RFP labeled cells were resuspended in Matrigel to restricted next to the brain slice. (d) Brain slice invasion assay. Left panel: Representative confocal images of each condition. Yellow dash lines indicate the border of brain slices. Brain slices locate on the left of the dash lines. Right panel: Quantification of invading colony numbers and invaded distance into brain slice using Image J. EV; empty vector transfected cells. RET-OE; RET overexpressing cells. shNTC; cells transfected with non-targeting control shRNA. shRET; cells transfected with shRNA targeting hRET. ****, p < 0.0001. ns, no significance.

In order to examine the effects of RET overexpression on invasion of brain tissue, we utilized an organotypic co-culture system of Matrigel-embedded breast cancer cells and mouse brain slices, previously described by Chuang et. al [38] in order to more closely mimic the physiological conditions underlying breast cancer cell invasion of the brain parenchyma. Cells were resuspended in a Matrigel spheroid, which was maintained immediately adjacent to the brain slice for five days of co-culture (Fig. 4c). RET overexpression significantly increased colonization and invasion of co-cultured brain slices, which were quantified using invaded distance and colony number within brain slice imaging fields respectively (Fig. 4d). RET knockdown resulted in reduced invasion relative to non-targeting control, however, this difference was not statistically significant. In sum, these data indicate a requirement of RET expression to facilitate invasion and colonization of brain tissue, concurrent with the in vitro observation of GDNF-increased metastatic capabilities. When considered alongside the effects of GDNF on in vitro invasion and chemotaxis, our *ex vivo* data suggest that brain-derived GDNF may act as an attractive factor for RET-overexpressing breast cancer cells, thereby enhancing tissue colonization.

Taken together, these findings lead to the conclusion that components of the brain microenvironment specifically activate RET signaling, and thus enhance both invasion and colony formation by breast cancer cells in *ex vivo* brain tissue. Considering the GDNF-RET dependent mechanism revealed by *in vitro* phenotypic assays, together with the relative abundance of the RET ligand GDNF in the brain relative to the liver, these results support the hypothesis that GDNF in the brain microenvironment facilitates colonization, invasion, and migration of RET-overexpressing breast cancer cells, thereby promoting dissemination of these cells in the brain.

### RET overexpression enhances brain colonization of MCF-7 cells *in vivo*

To investigate whether RET overexpression facilitates formation of breast cancer BrM *in vivo*, we established NSG mouse xenografts through intracardiac injection of RFP-luciferase labeled MCF-7 RET-OE cells. Successful injection of breast cancer cells was confirmed by disseminated luminescence immediately after cell injection (Fig. S4a). At 7 weeks post-injection, tumor growth was increased in RET-OE xenografts compared to EV controls, reflected by a significant difference in area under the curve (AUC) of bioluminescence relative to week 0 (Fig. 5b). Further, fold change in brain luminescence at the time of death or euthanasia was significantly higher in RET-OE xenografts than EV controls (Fig. 5c). Based on bioluminescence overlayed with x-ray images acquired 7 weeks post-injection, brain metastases developed in 100% of animals injected with RET-OE cells, compared to only 50% of the animals injected with EV cells. All brain metastases were confimred as RFP-positive (Fig. 5g). Overall survival of the mice was recorded with euthanasia performed when animal showed the IACUC pre-defined clinical endpoints. Xenografts implanted with RET-OE cells displayed reduced overall survival relative to the control group (p = 0.03) (Fig. 5e). Similar numbers of macro-metastases (macromets) to non-brain organs were identified in both RET-OE and EV groups, with RFP detected in the adrenal glands, ovaries, kidney, liver, and gallbladder (Fig. 5f-g). Diffuse RFP signatures were detected in the lungs of multiple animals, but no lung macromets were observed. These observations indicate that high levels of RET expression facilitate brain-tropic colonization of breast cancer cells, which is associated with poorer survival in NSG mice.

**Figure 5.**
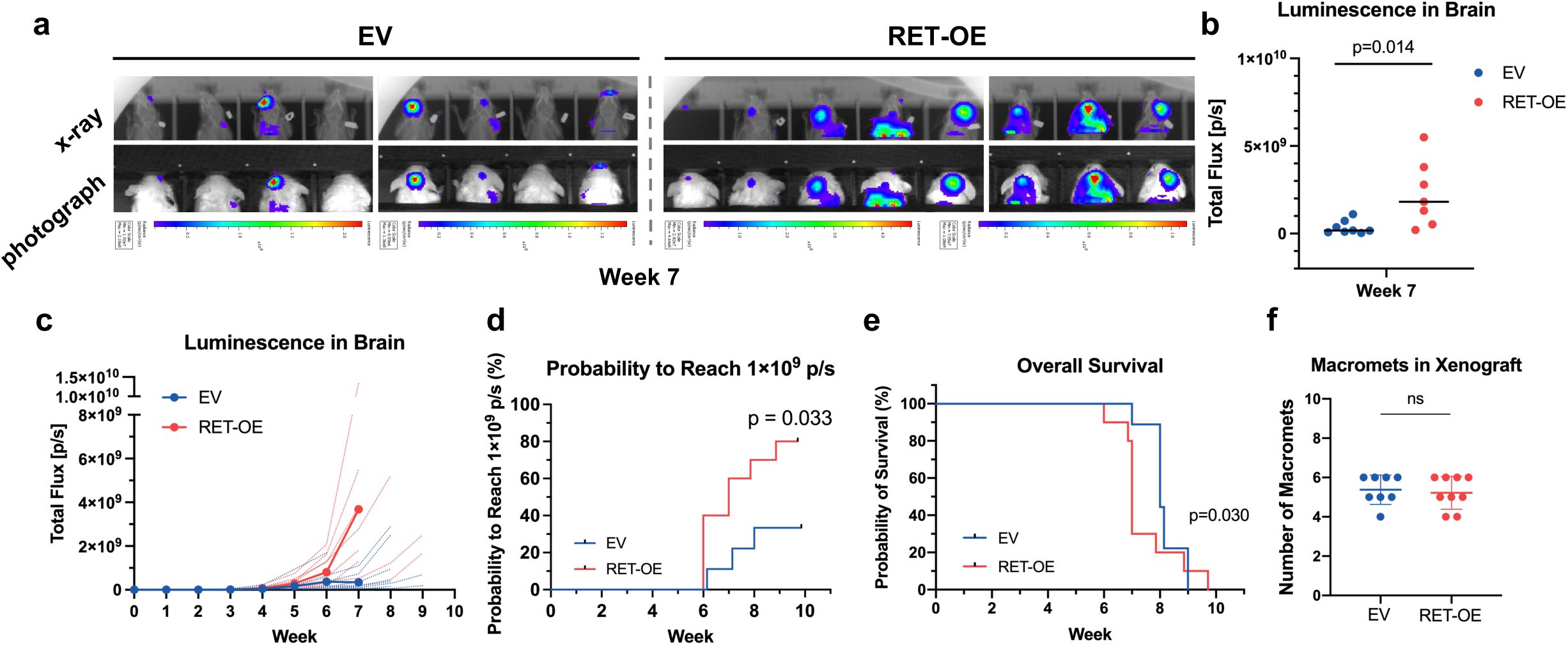
RET overexpression enhances brain colonization of MCF-7 cells *in vivo*. (a) Bioluminescence overlapped with x-ray images (top panel) or photographs (bottom panel) of the individual animals alive at week 7 (8/9 in EV and 8/10 in RET-OE) post intra-cardiac injection. All animals in RET-OE group showed concentrated luminescence in brain that indicates BrM formation. (b-c) Quantification of luminescence in brain in week 7 (b) and throughout the whole experiment (c). Individual reads were implicated by dash lines. Averages from week 8 were removed from the plot due to animal death. One outlier was removed for unpaired t test in (b). (d) Probability of luminescence in brain to reach 1×10^9 p/s. Breslow-Wilcoxon test. (e) Overall survival. (f) Numbers of macromats counted after animal euthanasia. EV; animals injected with RFP-labeled empty vector transfected MCF-7 cells. RET-OE; animals injected with RFP-labeled RET overexpressing cells.

To investigate the role of RET signaling in lobular breast cancer brain metastasis (BrM) progression, we examined the effect of knockdown of RET using shRNA in the SUM44 cell line. NSG mice were injected with SUM44 cells via the mammary fat pad. Survival analysis indicated that the shRET group exhibited a longer median survival compared to the shNTC group (Fig. S5a). Brain metastases formed in 6/7 mice in the shNTC group and 5/6 mice in the shRET group (Fig S5b, Table S1).The first detectable brain metastasis signal was set as the baseline. Brain metastasis progression was assessed at early (week 5) and late (week 10) time points using IVIS imaging (Fig. S5c-d). One mouse from each group was excluded from the analysis due to the lack of brain metastasis. At both early and late time points, the shNTC group consistently demonstrated a greater fold increase in brain metastasis signal intensity compared to the shRET group, suggesting more rapid metastatic growth in the presence of RET. Although we initially started with 10 mice in each group, the sample size decreased over the course of the experiment due to complications during the primary tumor resection and animal deaths, which prevented tissue harvesting. While reduced sample size lowered the statistical power and significance, the observed trends suggest a potential role for RET in facilitating the growth of lobular breast cancer brain metastases.

### CREB or P70S6K knockdown attenuates GDNF-induced RET signaling and cell migration

To further explore the mechanism of BrM development mediated by RET signaling, we employed a Human Phospho-Kinase Array (R&D), which facilitates screening of relative phosphorylation levels of 43 kinase phosphorylation sites, along with total expression of 2 proteins involved in the RET signaling network. Increased levels of phosphorylated targets such as P70-S6 Kinase (P70S6K), ERK1/2, AKT and STAT3 were observed in GDNF-stimulated empty vector control cells, and this effect was markedly increased in GDNF-treated RET-OE cells (Fig. 6a). Interestingly, the phosphorylation of many Src kinase family members (e.g. Lyn, Src, Fyn) was downregulated by GDNF. To gain a comprehensive view of GDNF-mediated RET signaling in breast cancer, we next performed reverse phase proteomic analysis (RPPA) on GDNF-treated sample pools in both MCF-7 (EV and RET-OE) and SUM44 (shNTC and shRET) models. Overall, there were 133 significant (p<0.05) changes in phosphorylation observed in the MCF-7 samples (Fig. S4b). The 40 targets exhibiting the highest fold change in MCF-7 RET-OE samples are shown (Fig. 6b). RPPA data were similar to the phospho-kinase array, with RET overexpression enhancing the GDNF-responsiveness of numerous signaling molecules. Specifically, increased phosphorylation of ERK (MAPK), AKT, P70S6K, and CREB were well-conserved from the phospho-kinase array, and ranked among the top 5 by logFC in GDNF-treated RET overexpressing cells in both characterization strategies. RPPA also revealed a significant increase in PLCγ-1 activation in response of GDNF stimulation after RET overexpression. Consistent with our observations in RET-overexpressing MCF-7 cells, activation of ERK, CREB, and PLCγ-1 was reduced by RET knockdown in the SUM44 cell line (Fig. 6b). Increased activation of the proteins of interest was confirmed in RET-OE cells via immunoblot. Altogether, identification of these downstream targets suggests possible molecular mechanisms of GDNF-RET pathway-mediated tumor progression.

**Figure 6.**
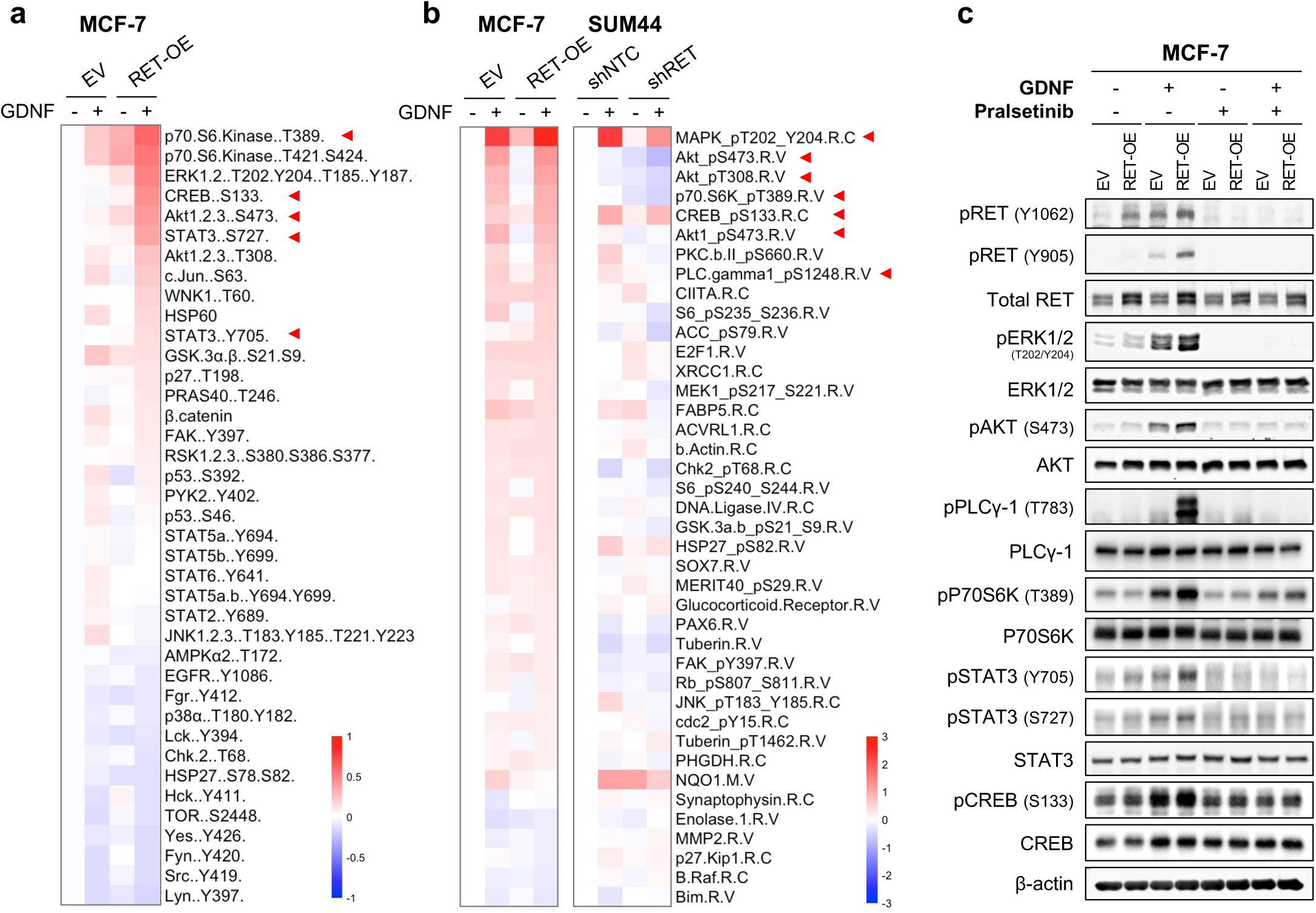
Phospho-kinase array and RPPA reveal increased activation of ERK1/2, AKT, PLCγ-1, P70S6K, STAT3, and CREB in RET overexpressing MCF-7 cells. (a) Normalized logFC of (phospho)-protein levels relative to the control group in Human Phospho-Kinase Array. All targets with significant (adjusted p value < 0.05) differences were plotted and sorted by logFC in RET-OE + GDNF group. (b) Normalized logFC of (phospho)-protein levels relative to the control group in MCF-7 or SUM44, respectively, in RPPA. Top 40 significant proteins were plotted and sorted by log FC in MCF-7 RET-OE + GDNF group. (c) Immunoblotting in RET overexpressing cell line model. ERK1/2, AKT, P70S6K, PLCγ, STAT3 and CREB are GDNF-responsive RET downstream targets that can be inhibited by treatment of Pralsetinib. Specificity of RET pathway signaling was confirmed by shRNA knockdown. GDNF, 100 ng/ml, 30 min. Pralsetinib, 500 nM. Vehicle, distilled water for GDNF, and DMSO for Pralsetinib. EV; empty vector transfected cells. RET-OE; RET overexpressing cells. shNTC; cells transfected with non-targeting control shRNA. shRET; cells transfected with shRNA targeting hRET.

To investigate whether the identified targets are involved in GDNF-stimulated pro-metastatic cell behaviors, we used siRNA to decrease expression of CREB, P70S6K, and PLCγ-1. Immunoblotting confirmed the efficiency of CREB, P70S6K, and PLCγ-1 knockdown in MCF-7 EV and RET-OE models (Fig. 7a-b, Fig. S7a). GDNF-induced phosphorylation of RET, ERK, AKT, and PLCγ-1 was attenuated with either CREB or P70S6K knockdown. Furthermore, using Boyden chamber migration assays, we observed that knockdown of CREB and P70S6K attenuates GDNF-stimulated enhancement of cell migration in both EV and RET-OE cell lines. However, there was no significant difference of GDNF-promoted cell migration between the groups with and without knockdown of PLCγ-1. Taken together, these data suggest that CREB and P70S6K may represent the mechanism by which GDNF activates cellular motility in RET-expressing breast cancer cells.

**Figure 7.**
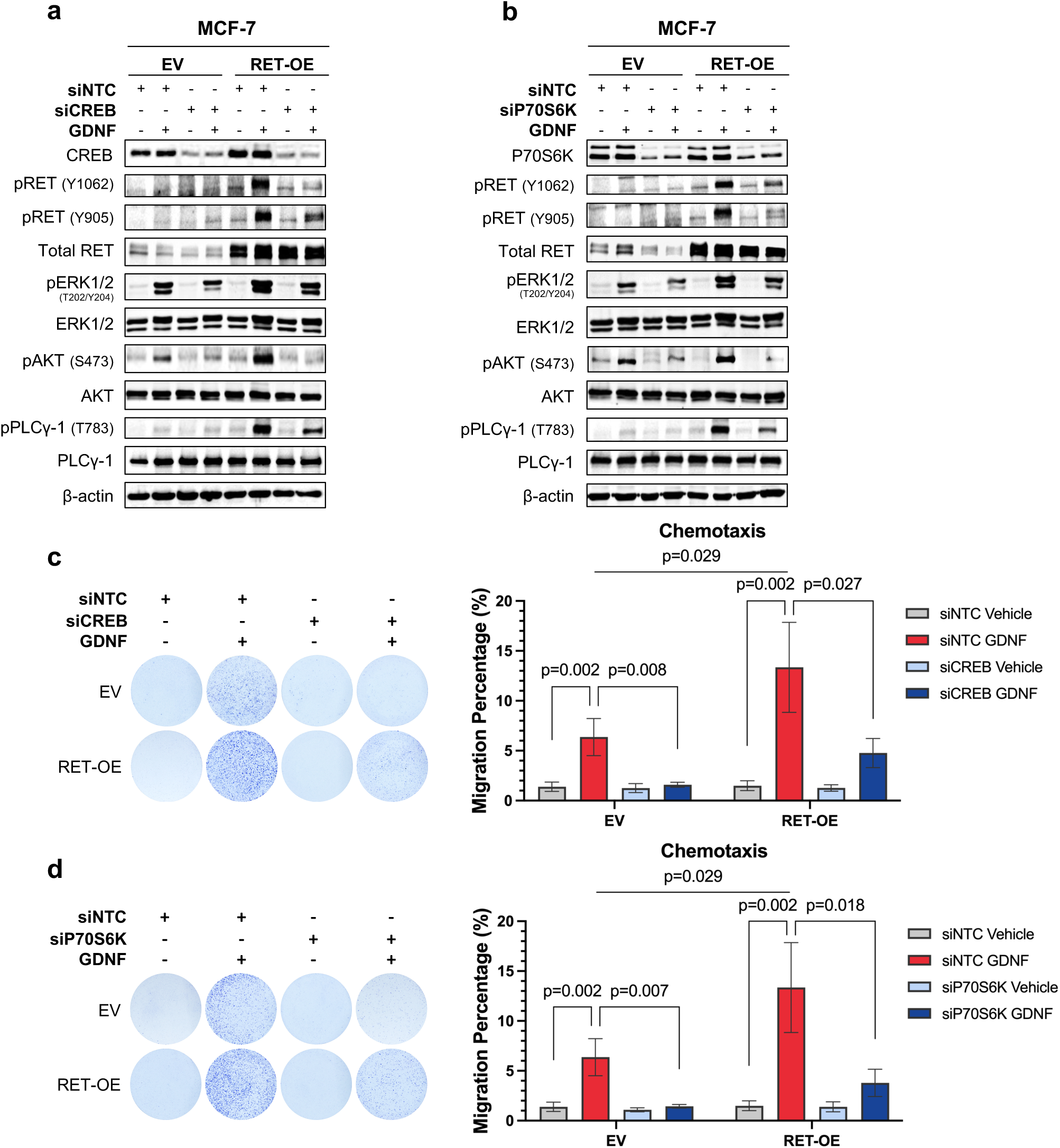
CREB or P70S6K knockdown attenuates GDNF stimulated activation of RET downstream targets and enhancement of cell migration. (a-b). immunoblotting in empty vector or RET overexpressing MCF-7 cell lines. ERK1/2, AKT, and PLCγ-1 are GDNF-responsive RET downstream targets that can be downregulated by CREB (a) or P70S6K (b) knockdown. (c-d) images and quantification of crystal violet–stained transwell inserts showing the migration of empty vector or RET overexpressing MCF-7 cells transfected with siCREB, siP70S6K or non-targeting control RNA in response to GDNF chemotaxis. One way ANOVA was performed to test for statistical significance in c & d. GDNF, 100 ng/ml, 30 and vehicle is distilled water for GDNF. EV; empty vector. RET OE; RET overexpressing cells. NTC; non targeting control RNA.

Consistent with our observations in RET-overexpressing MCF-7 cells, activation of ERK, CREB, and PLCγ-1 was reduced by RET knockdown in the SUM44 cell line (Fig. 6b). Increased activation of the proteins of interest was confirmed in RET-OE cells via immunoblot. Altogether, identification of these downstream targets suggests possible molecular mechanisms of GDNF-RET pathway-mediated tumor progression.

## Discussion

The diagnosis of brain metastasis poses a remarkable clinical challenge to breast cancer patients [46]. The reported incidence of BrM from breast cancer is rising due to early identification of BrM lesions and increasingly effective control of extracranial metastatic disease, which extend survival but increase lifetime risk of developing BrM [47]. The current understanding of breast cancer BrM is mostly based on studies of TNBC and HER2-enriched breast cancers, for which multiple systemic therapies are now available, including HER2-targeting tyrosine kinase inhibitors and trastuzumab conjugates [48–51]. In contrast, specific drivers of BrM from ER+BC have not been well-characterized, leaving patients experiencing this complication of the most common breast cancer subtype with few therapeutic options. We previously reported RET, a receptor tyrosine kinase corresponding to a brain-specific growth factor key to brain development and glial cell protection (GDNF), as one of the most frequently enriched clinically actionable genes in patient-matched brain metastases relative to primary tumors, with gains in expression in 38% of breast cancer BrM samples, with a pronounced enrichment in patients who diagnosed with ER+ primary breast cancers [4]. Given that GDNF is abundantly and specifically expressed in the brain microenvironment, we hypothesized that the availability of GDNF in the brain provides a permissive microenvironment for RET-overexpressing breast cancer cells to form brain metastases, and that RET inhibition is a potential therapeutic strategy for breast cancer brain metastases driven by elevated RET expression.

Our study extends the findings of a previous study [6] that demonstrated the role of RET overexpression in promoting cancer cell adhesion, invasion and outgrowth in the brain, where EGFR was required to deliver a pro-brain metastatic phenotype. While the earlier study focused on BDNF, our work investigates the effects of GDNF, another neurotrophic factor, on RET-mediated brain colonization. We incorporated orthotopic mouse models, providing in vivo evidence that RET overexpression significantly enhances brain metastatic potential in a more physiologically relevant way. Furthermore, our research goes a step further by exploring the downstream signaling pathways activated by RET. This detailed mechanistic analysis offers new insights into how RET contributes to brain metastasis through its signaling networks, which could indicate therapeutic strategies targeting RET in breast cancer patients with brain metastases. Thus, our study not only reinforces but also extends the current understanding of RET’s role in brain metastatic progression.

In this study, we characterized RET as a driver of breast cancer brain metastasis originating from ER+BC. Both RET expression and activation were upregulated in breast cancer BrM relative to the matched primary tumor, and relative to lesions from other sites of metastasis. Elevated RET expression and GDNF-RGS score were associated with poor prognosis in ER+BC patients, supporting the prognostic significance of RET. These data are consistent with past reports of the significance of GDNF-RGS, which predicts poor prognosis in other three ER+ breast cancer cohorts (NKI295, Pawitan, TransBig) [26], which highlights the predictive power of gene signatures, rather than single gene expression, to assess prognosis of ER+BC.

Using cell line models representative of the two most common histologic subtypes of breast cancer, we found that activation of RET, along with downstream ERK and AKT was upregulated in RET overexpressing cells in response to GDNF, which was potently inhibited by Pralsetinib treatment. The specificity of GDNF-RET signaling was confirmed through shRNA-mediated RET knockdown. We observed a time-dependent increase in the level of RET phosphorylation, which peaked at different time points in MCF-7 and SUM44 cells after GDNF treatment. These findings highlight the dynamic nature of RET activation. Further, we confirmed a role for RET overexpression and activation in the mediation of pro-metastatic properties *in vitro*, such as migration, invasion and anchorage-dependent growth. They were mediated by GDNF, which serves as a chemoattractant for RET-overexpressing cells. Additionally, RET overexpression alone does not facilitate cell growth, further highlighting the critical role of ligand availability in RET-driven breast cancer phenotypes. This GDNF-induced migratory ability is abrogated by administration of Pralsetinib, indicating its potential as a therapeutic intervention. Furthermore, brain organotypic co-culture experiments revealed that RET overexpression specifically promoted brain tissue colonization and invasion compared to liver, while Pralsetinib treatment significantly reduced these effects.

In vivo studies provided additional support for RET’s role in brain metastases. RET overexpression was associated with earlier BrM formation, increased brain luminescence, and shorter survival in NSG mice. However, the absolute number of macrometastases was comparable between RET-overexpressing and control groups, suggesting that RET may enhance colonization efficiency and metastatic growth rather than overall tumor seeding. These findings reinforce the critical role of RET signaling in BrM progression and suggest therapeutic targeting of RET as a strategy to delay or prevent brain metastases.

At signaling level, Human Phospho-Kinase Array and RPPA identified numerous signaling molecules responsive to GDNF-RET activation in the selected breast cancer cell models, including ERK1/2, AKT, PLCγ, P70S6K, STAT3 and CREB, which are associated with malignant phenotypes and oncogenic processes including cell survival, proliferation, migration, and invasion. Our study found that either CREB or P70S6K knockdown attenuates GDNF-stimulated activation of RET downstream targets and resulting enhancement of cell migration. PI3K/AKT and ERK/MAPK pathways are involved in P70S6K and CREB regulation in previous literatures. CREB activation via PI3K/AKT and ERK/MAPK pathways mediates cell proliferation and motility in prostate cancers [52]. Pre-treatment with specific AKT and ERK inhibitors reduces activation of CREB [53]. Moreover, inhibition of either PI3K or ERK induces prompt and prolonged suppression of p70S6K activation [54]. However, decreased activation of AKT and ERK after knockdown of P70S6K or CREB has not been reported. Therefore, we suspect a potential positive feedback loop between P70S6K, CREB, ERK, and AKT. Activation of PLCγ-1 occurs when cells experience stress or oxidative damage [43]. Overexpression of PLCγ-1 has been reported in various cancers including colorectal cancer and head and neck cancer, though the oncogenic mechanism remains undepicted [44, 45]. However, despite GDNF-induced phosphorylation of RET and ERK was reduced, AKT remained activated after PLCγ-1 knockdown. Furthermore, treatment of siRNA targeting *PLCG1* exhibited no effect on GDNF-promoted cell migration of RET overexpressing breast cancer cells. Inhibition of AKT significantly reduced cell migration in breast and gallbladder cancer cells [55, 56]. Collectively, these findings suggest that GDNF promotes cell migration in RET overexpressing breast cancer cells through crosstalk between the AKT/ P70S6K/CREB signaling pathways.

Overall, our findings support the hypothesis RET overexpression increases metastatic potential of ER+BC cells with a specific increase in colonization of brain tissue, the principal site of RET ligand expression. We show that Pralsetinib is a potent inhibitor of RET’s functions in breast cancer cells in *in vitro* and *ex vivo* assays. Critical future directions of research include validation of the activity RET-selective inhibitors in *in vivo* models of breast cancer brain metastasis. Additionally, our proteomic studies provide a description of GDNF-stimulated RET signaling networks for further follow-up studies. Other recent studies [6] have identified novel signaling partners for RET in the setting of BC BrM, including EGFR and NTRK2. Given the known relationship between RET and TRK overexpression in other cancers of the central nervous system [57] and the availability of potent dual RET-TRK small molecule inhibitors [58], clarification of the relationship between RET, TRK receptors, and the downstream pathways identified in this study warrant further investigation. Additionally, RET expression has recently been shown to be upregulated in the setting of *ESR1* fusions [59], and we have previously shown that ESR1 mutations are enriched in samples of BrM from ER+BC [60]. Taken with recent studies highlighting the role of RET in BrM, signaling interactions between RET and altered ER in the setting of brain-metastatic disease may be a valuable future direction of research.

In summary, our study provides insight into the role of RET in the formation of breast cancer brain metastases, and provides initial characterization of RET-selective inhibitors in ER+ breast cancer Considering the recent FDA approval of Pralsetinib and Selpercatinib for use in MTC, our findings show strong potential for clinical translation.

## Materials and Methods

Additional details are provided in the Supplementary Materials and Methods section.

### RNA Sequencing

Sample collection and processing for paired bone and brain metastasis were described in the previous literature [4, 61]. Collection for ovary and GI metastasis cohort was similarly processed. Quality control was performed by FastQC (https://www.bioinformatics.babraham.ac.uk/projects/fastqc/) and summarized with multiQC (https://multiqc.info/). RNA expression was quantified with Salmon[62] at quasi-mapping mode, and normalized to log2 trimmed mean counts per million (log2TMMcpm). For gene set scoring, Gene set variation analysis (GSVA) [63] was performed on normalized data with selected gene sets as specified in individual sections.

Besides public RNA or scRNAseq data specified above, this study also used TCGA and METABRIC in analysis. METABRIC RNA microarray data was downloaded from Synapse (RRID:SCR_006307) and log normalized[64]. GSVA of selected gene set was performed with GSVA R package, in ssgsea or gsva mode with default parameters [63, 65].

For survival analysis, only estrogen receptor positive (ER+) and LumA patients were selected to avoid influence caused alone by PAM50 subtype. DormSig ssGSEA scores were stratified into low and high levels with optimized threshold using in surv_cutpoint and surv_categorize implemented in R survminer package (https://cran.r-project.org/web/packages/survminer/index.html). Kaplan-Meier curve were plotted for each group using disease free survival, with p value from log-rank test, and hazard ratio (HR) calculated using univariate Cox regression (coxph in R survival package, https://cran.r-project.org/web/packages/survival/index.html).

### Tissue Culture and Cell Model Generation

MCF-7 (RRID: CVCL_0031), T-47D (RRID: CVCL_0553), MDA-MB-134-VI (MM134, RRID: CVCL_0617), and HEK293T (293T, RRID: CVCL_0063) were purchased from American Type Culture Collection (ATCC). SUM44PE cells (SUM44, CVCL_3424) were purchased from Asterand. Cell identity was authenticated by Short Tandem Repeat DNA profiling (University of Arizona), and cells were routinely tested to be mycoplasma free.

To establish RET overexpressing (RET-OE) cell line models in MCF-7, MM134 and T-47D, a lentiviral vector carrying RET or paired empty vector control (VectorBuilder) was first packaged in HEK293T cells to generate viral particles by PEI transfection with pMDL, pRev and pVSVG. Supernatant was collected at 48 hr, 60 hr and 72 hr post-transfection to infect the target breast cancer cells followed by blasticidin selection.

For stable RET deficient (shRET) breast cancer cell line model development, 3 shRNAs with distinct sequences to target RET were designed (Dharmacon) with one non-targeting control shRNA provided (Dharmacon). Lentiviral shRNA was packaged with psPAX2 and pMD2.G vectors in HEK293T cells, and virus-containing supernatant was collected at 48 hr, 60 hr and 72 hr post-transfection to infect the target breast cancer cells (MCF-7 and SUM44PE) followed by puromycin selection.

To label the established cell line models with tRFP and luciferase, a lentiviral vector was designed that carries both turbo-RFP and luciferase. The same lentivirus packaging procedures was followed as described above in shRNA transfection. The lentivirus-infected target cells were selected for 2 weeks with optimized concentration of G418 (400 µg/ml for MCF-7 RET overexpression and RET knockdown models as well as SUM44 RET knockdown models). Cells with top 15% RFP expression level were gated and collected from Fluorescence-Activated Cell Sorting (FACS, BD Biosciences) and recovered in G418 selection medium added with 1X Penicillin-Streptomycin for an extra week before back into regular culture medium.

### Human Phospho-Kinase Array Analysis

The commercial Proteome Profiler Human Phospho-Kinase Array Kit (R&D Systems) was employed for Phospho-Kinase Array. Cells were plated in 1 10 cm dish per condition and serum-starved in 0.5% FBS medium (0.5% CSS medium for SUM44PE) for 18 hours, followed by 30 min treatment of 100 ng/ml GDNF or vehicle (distilled water) in 0.5% FBS medium (0.5% CSS medium for SUM44PE). Whole cell lysates were prepared following the instruction provided by the manufacturer. Protein concentrations were determined by Bicinchoninic Acid (BCA) Protein Assay Kit (Pierce™) and 600 ug protein was calculated accordingly to incubate with each membrane. The blotting and development procedures were also completed following the manufacturer’s handbook. Finally, the developed blots were exposed to films for 1, 2, 5, 8 and 10 min followed by processing using Konica SRX-101A Film Processor, and 8-min exposure was optimized for quantification. Films were scanned under colorimetric-gray mode in ChemiDoc XRS+ Imaging System (Bio-Rad) and the .scn files were analyzed using Image Lab Software (Bio-Rad). Background of the blots was substracted and integrated density (pixel) of each dot on the blots was quantified through Image Lab - Volume Tools.

For data processing, relative protein levels were determined by normalization to the integrated density of PBS spots provided by the manufacturer on the blot as negative control (calculated as “0”) and to the conserved reference spots as positive control (calculated as “1”). Comparison between different treatment groups and the control group (vehicle-treated empty vector (EV, GDNF-) group) was accomplished by using limma package in R, with results sorted by adjusted p value. All 39 significant (adjusted p value < 0.05) targets were plotted into heatmap ranked by LogFC in RET-OE + GDNF group. Heatmaps of all significant targets in hierarchical clustering analysis could be found in Figure S6 [66].

### Reverse Phase Protein Array (RPPA)

Before treatment, cells were incubated in 0.5% FBS medium (0.5% CSS medium for SUM44PE) for 18 hours, and then switch to 100 ng/ml GDNF or vehicle (distilled water) in 0.5% FBS medium (0.5% CSS medium for SUM44PE) for 30 min. Cell lysate was prepared following a protocol modified from that provided by MD Anderson Center. Cells were harvested in RPPA lysis buffer with protease inhibitors. Protein concentrations were determined by BCA Protein Assay Kit (Pierce™). Then, lysates were mixed with RPPA lysis buffer and 4X SDS/B-Me sample buffer (40% Glycerol, 8% SDS, 0.25 M Tris-HCL, pH 6.8 and Beta-mercaptoethanol (B-Me) added before use at 1/10) to have final concentration of 1.22 μg/μl for further process by MD Anderson Center. Cell lysate samples were serially diluted two-fold for 5 dilutions (undiluted, 1:2, 1:4, 1:8; 1:16) and arrayed on nitrocellulose-coated slides in an 11×11 format to produce sample spots. Sample spots were then probed with antibodies by a tyramide-based signal amplification approach and visualized by DAB colorimetric reaction to produce stained slides. Stained slides were scanned on a Huron TissueScope scanner to produce 16-bit tiff images. Sample spots in tiff images were identified and their densities quantified by Array-Pro Analyzer.

For data processing, comparison between different treatment groups and the corresponding control group (vehicle-treated empty vector (EV, GDNF-) group in MCF-7 or vehicle-treated non-targeting control (shNTC, GDNF-) group in SUM44 models, respectively) was accomplished by limma package in R, with results sorted by adjusted p value (Table S2). Top 40 significant (adjusted p value < 0.05) targets were selected and plotted into heatmap ranked by LogFC in RET-OE + GDNF group (shNTC + GDNF group in SUM44). Heatmaps of all significant targets in hierarchical clustering analysis could be found in Figure S6.

### Western Blotting (WB)

Cells were all pre-incubated in low serum medium (0.5% FBS medium for MCF-7, T-47D and MM134 and 0.5% CSS medium for SUM44) for 16 hr (serum starvation) followed by 30 min of GDNF treatment. Pralsetinib (Blueprint Medicine, GAVRETO™) was provided not only in the low serum medium during serum starvation, but also applied during GDNF (or distilled water) treatment, DMSO as the vehicle control. Specially, 10 ng/ml GFRα1 was applied together with GDNF for T-47D. Cell lysates were harvested in RIPA buffer with Halt Protease and Phosphatase Inhibitor Single-Use Cocktail added. Protein concentrations were determined by BCA Protein Assay Kit. 50 µg protein was loated to each well of a 15-well 8% denaturing gel. Both Li-Cor Odyssey Western Blotting protocol and Bio-Rad General Western Blotting protocol were followed as adaptive to Odyssey® Imaging System (Li-Cor) and ChemiDoc XRS+ Imaging System (Bio-Rad), respectively. For quantification, the intensity (pixel) of the bands was measured by Image J. Relative abundance was calculated by normalization to protein level of β-actin, and intensity of the target in GDNF-treated group without application of the inhibitor was set as 100%.

### Cell Growth Assay

Cells were serum-starved in 0.5% FBS medium overnight before seeded in 96-well flat bottom tissue culture plates for 2D level study or Corning® 96 Well Clear Flat Bottom Ultra Low Attachment Microplates for ULA growth assay, both at 3,000 cells per well in 60 ul 0.5% FBS medium. 24 hr after cell seeding, cells were treated with drugs in extra 40 ul 0.5% FBS DMEM to reach a final concentration of 100 ng/ml GDNF (distilled water as vehicle) and 500 nM Pralsetinib (DMSO as vehicle, ATCC®). For GDNF dose-response assay, 9 different concentrations of the drug were prepared by 1:2 serial dilution beginning at 100 ng/ml, with distilled water as vehicle. ATP levels in the culture medium were determined using a CellTiter-Glo® Luminescent Cell Viability Assay to reflect cell proliferation. A protocol provided by the manufacturer was followed. Luminescence was acquired using GloMax® Microplate Reader or DTX 880 Multimode Detecter. For data processing, luminescence of each well was corrected with the reads of the blank (0.5% FBS medium without any cell). All reads were normalized to Day 0 plates and fold change relative to Day 0 was used to reflect cell growth.

### Dose Response Assay

MCF-7 cells were seeded as described above. For drug preparation, 9 different concentrations of Pralsetinib were prepared by 1:5 serial dilution of a 4 mM solution. ATP levels in the culture medium were measured following the same protocol discussed in the previous section. Nonlinear regression and statistical analysis were performed in Graphpad Prism.

### Sphere Formation Assay

MCF-7 cells were plated in Corning® 96 Well Clear Round Bottom Ultra Low Attachment Microplate at 5,000 cells in 50 ul 0.5% FBS DMEM per well. After overnight incubation, cells were treated with additional 50 ul medium to reach a final concentration of 100 ng/ml for GDNF and 100 nM for Pralsetinib, with distilled water and DMSO as the vehicle, respectively. Spheres formed by MCF-7 cells were imaged by Olympus IX83 Microscope (4X) after 7-day culture. The diameter of each sphere was determined by Image J, and sphere area was calculated accordingly.

### Soft Agar Assay

The protocol was modified from a previous publication[37]. To generate the agar layers, 0.6% agar was prepared by 1:1 mixture of 1.2% Agar stock (1.2 g Agar in 100 ml distilled water) and enhanced medium (DMEM plus 20% FBS, with 1X NEAA (Thermo Scientific) and 1X Antibiotic-Antimycotic (Thermo Scientific). After solidification of the agar layer, cells resuspended in 0.4% agar in enhanced media were seeded at 10,000 cells per well in 6-well plate. After overnight incubation, GDNF (100 ng/ml) and/or Pralsetinib (100 nM) were administrated. Cells were incubated for 7 weeks, after which the cells were fixed and stained in Crystal Violet solution. 4X images were taken through Olympus IX83 Microscope, from which colony number and colony size were quantified in Image J.

### Wound Scratch Assay

Cells were pre-incubated in 0.5% FBS DMEM overnight before seeded in Incucyte® ImageLock 96-well Plate, which was freshly coated with 50 ul 100 µg/ml Growth Factor Reduced Matrigel (Corning) diluted in regular culture medium. 1.5×105 cells were seeded per well in 8 replicates. MCF-7 cells formed 100% confluent cell monolayer 24 hr after cell seeding. Medium was replaced with 100 ul DPBS before wound scratching. Incucyte® WoundMaker™ was employed to create homogeneous wounds in the middle of cell monolayer at a width of 700-800 μm. Then cells were treated with 100 ng/ml GDNF and 500 nM Pralsetinib or vehicle (DMSO) in 0.5% FBS DMEM. Mitomycin C (Sigma-Aldrich) was applied to prohibit cell proliferation at 5 ug/ml [67]. Cell migration was monitored by IncuCyte ZOOM® Live-Cell Analysis System with 10X images taken every 4 hr. Relative wound density was quantified in built-in Incucyte wound scratch module, reflecting portion of region healed by migrated cells to initial wound.

### Boyden Chamber Transwell Assay

Cells were pre-incubated in 0.5% FBS DMEM overnight before seeded in Transparent PET membrane 8 μm pore size insert (Thermo Fisher Scientific). 500,000 cells in 100 ul 0.5% FBS DMEM were seeded per insert in 24-well plates. For transwell invasion assay, both sides of the inserts were coated with 25 μg/ml Collagen I, Rat Tail (Corning) cross-linked in 0.02N acetic acid. For both chemotaxis and invasion assays, 100 ng/ml GDNF or vehicle (distilled water) in 700 ul 0.5% FBS DMEM was added to the bottom chamber, while the inhibitor Pralsetinib (500 nM) was applied to the upper chamber (insert) where the cells were seeded. Cells were incubated for 48 hours, after which the excess cells in top chambers were removed and cells attached to the bottom side of the inserts were fixed and stained in Crystal Violet solution (10% Methanol, 1% Paraformaldehyde in PBS and 0.01% Crystal Violet,). 1X images were taken through Olympus SZX16 dissecting microscope, from which migration percentage was quantified under Image J – Area Fraction to reflect the area taken by purple-stained migrated cells divided by the area of whole insert.

### Brain Slice Colony Formation Assay

Brains were obtained from 6-8-week-old mice. Brain slices were sectioned at 300 micron using a vibratome generously provided by Dr. Gerard Apodaca’s lab. Sections were placed on a Millicell Cell Culture Insert (30 mm, hydrophilic PTFE, 0.4 µm, Millipore Sigma) and cultured overnight for recovery before cell addition. After 24 hours, 50,000 RFP-labeled MCF-7 cells were seeded on the slices restricted by a cylinder and cultured for 5 days before tissues were gently washed in PBS (with calcium and magnesium) for 5 minutes, followed by RFP channel imaging (Olympus IX83 Microscope, 4X).

Quantification of cell number in each group was completed in Image J. Each dataset represents n=36 samples scored between 3 independent experiments. Unpaired t-test was performed for statistical analysis.

### Brain Slice Invasion Assay

Brain slices were generated as illustrated above and placed on the Millicell Cell Culture Insert. After overnight recovery, 100,000 RFP-labeled RET overexpressing MCF-7 cells were resuspended in Growth Factor Reduced Matrigel (Corning) and the Matrigel spheroid was dropped next to the brain slice (Fig. 4c). After 5-day co-culture, the brain slices were fixed in 10% formalin for Z-stack confocal microscope imaging (Nikon A1R Confocal Microscope, 10X). Quantification was completed in a double-blind manner. Cell colonies invaded into brain slices was quantified using multipoint tool in Image J. For invaded distance, the distance (pixel) from the farthest invaded cell to tangent line of the edge of brain lice was measured in Image J. Unpaired t-test was used for statistical analysis.

### *In vivo* Xenograft Study

Four-week-old female NSG immunocompromised mice were ordered from the Jackson Laboratory (#005557, NOD/NSG) according to University of Pittsburgh IACUC approved protocol #21028901. Animals were implanted with slow-release Estradiol (E2) wax pellet (0.5 mg) disinfected by radiation one day before injection, and were dosed with ketofen at 5 mg/kg once daily beginning at the date of surgery for 3 continuous days. For intra-cardiac injection, RFP-luciferase labeled MCF-7 RET over-expressing cells and the empty vector control were resuspended in PBS at 1,000,000 cells/ml. 100 μl cell suspension (100,000 cells) was then injected into left ventricle of NSG mice with 9 and 10 animals in EV or RET-OE group, respectively. The intra-cardiac injection was operated by Mrs. Christy Smolak. Successful injection was confirmed by intraperitoneal injection of D-Luciferin (150 mg/kg) and bioluminescence imaging (BLI) with IVIS® Lumina X5 Imaging System.

IVIS images of the mice were taken weekly. As level of E2 released by the wax pellet dropped, 8 ug/ml Estradiol was provided in the water from the 7^th^ week post injection. According to the IACUC protocol, mice were euthanized when the following pre-defined signs of euthanasia were observed: above 20% weight loss, difficulty ambulating, anorexia, piloerection, hunched posture, rough and ungroomed hair coat and ungroomed appearance, excessive scratching and licking, mutilation of painful area, pallor due to severe anemia, tumor size > 1600 mm3, respiratory distress, and not responsive to external stimuli. Animal survival was recorded. Macro-mets as well as other potential organs (lung, liver and UG tracts) for metastatic spread were harvested and imaged under Olympus SZX16 Microscope to detect RFP expressed by the implanted tumor cells.

## Acknowledgements

This study was in part supported by the Breast Cancer Alliance and METAvivor and UPMC Hillman Cancer Center P30CAXXX. This work used the HTC cluster, which is supported by NIH award number S10OD028483 and the Animal Facility of the UPMC Hillman Cancer Center shared resource which are supported in part by award P30CA047904. The Functional Proteomics RPPA Core facility is supported by MD Anderson Cancer Center (Support Grant # 5 P30 CA016672-40). We appreciate technical and equipment support from Dr. Gerard Apodaca’s lab for vibratome training and usage. Simeng Liu, Ye Cao and Fangyuan Chen were visiting research scholars at the University of Pittsburgh School of Medicine supported by funds from The China Scholarship Council and Tsinghua University.

## Supplementary Materials and Methods

### Tissue Culture and Cell Model Generation

MCF-7 (RRID: CVCL_0031), T-47D (RRID: CVCL_0553), MDA-MB-134-VI (MM134, RRID: CVCL_0617), and HEK293T (293T, RRID: CVCL_0063) were purchased from American Type Culture Collection (ATCC). SUM44PE cells (SUM44, CVCL_3424) were purchased from Asterand. Cell identity was authenticated by Short Tandem Repeat DNA profiling (University of Arizona), and cells were routinely tested to be mycoplasma free. Cells were maintained in continuous culture for less than 3 months with medium as follows: MCF-7: DMEM, 10% fetal bovine serum; T-47D: RPMI-1640, 10% fetal bovine serum; MDA-MB-134-VI: DMEM:L15=1:1, 10% fetal bovine serum; SUM44PE: DMEM/F12, 2% charcoal stripped serum, 5 ug/ml insulin, 1 ug/ml hydrocortisone, 10 mM HEPES, 5 mM ethanolamine, 5 ug/ml transferrin, 10 nM triiodothyronine, 50 nM sodium selenite; HEK293T: DMEM, 10% fetal bovine serum.

To establish RET overexpressing (RET-OE) cell line models in MCF-7, MM134 and T-47D, a lentiviral vector carrying RET (pLV[Exp]-EGFP-T2A-Bsd-EF1A>hRET, VectorBuilder, Cat #VB190923-1153aty) or paired empty vector control (VectorBuilder, Cat #VB191024-2380bjs) was first packaged in HEK293T cells to generate viral particles by PEI transfection with pMDL, pRev and pVSVG. Supernatant was collected at 48 hr, 60 hr and 72 hr post-transfection to infect the target breast cancer cells followed by blasticidin (Thermo Fisher Science, Cat #A1113903) selection.

For stable RET deficient (shRET) breast cancer cell line model development, 3 shRNAs with distinct sequences to target RET were designed (Dharmacon) with one non-targeting control shRNA provided (Dharmacon). Lentiviral shRNA was packaged with psPAX2 and pMD2.G vectors in HEK293T cells, and virus-containing supernatant was collected at 48 hr, 60 hr and 72 hr post-transfection to infect the target breast cancer cells (MCF-7 and SUM44PE) followed by puromycin (Thermo Fisher Science Life Technology, Cat #A11138-03) selection.

To label the established cell line models with tRFP and luciferase, a lentiviral vector pLEX_TRC211-Luciferase2N-TurboRFP-c was designed that carries both turbo-RFP and luciferase. The same lentivirus packaging procedures was followed as described above in shRNA transfection. The lentivirus-infected target cells were selected for 2 weeks with optimized concentration of G418 (400 µg/ml for MCF-7 RET overexpression and RET knockdown models as well as SUM44 RET knockdown models). Cells with top 15% RFP expression level were gated and collected from Fluorescence-Activated Cell Sorting (FACS, BD FACSAria™ Fusion Cell Sorter, BD Biosciences) and recovered in G418 selection medium added with 1X Penicillin-Streptomycin (Hyclone, Cat #SV30010) for an extra week before back into regular culture medium.

### Reverse Phase Protein Array (RPPA)

Cells were plated in triplicates in 6-well plates, 2 wells per treatment. Before treatment, cells were incubated in 0.5% FBS medium (0.5% CSS medium for SUM44PE) for 18 hours, and then switch to 100 ng/ml GDNF or vehicle (distilled water) in 0.5% FBS medium (0.5% CSS medium for SUM44PE) for 30 min. Cell lysate was prepared following a protocol modified from that provided by MD Anderson Center. Cells were harvested on ice by washing in ice-cold PBS twice, and lysing immediately in 6-well plates in RPPA lysis buffer (1% TritonX-100, 50 mM HEPES, pH 7.4, 150 mM NaCl, 1.5 mM MgCl2, 1 mM EGTA, 100 mM NaF, 10 mM Na4P2O7, 1 mM Na3VO4, 10% Glycerol) with freshly added protease inhibitors (cOmplete™, EDTA-free Protease Inhibitor Cocktail, Roche, Cat #05056489001 and PhosSTOP™, Sigma-Aldrich, Cat #049068-37001). The plates were incubated on ice for 30 min and shaken every 5 min, followed by rocking at 4 °C for 10 min. Lysates were collected on ice into 1.5 ml eppendorf tube, incubated for 20 min and vortexed at 5 min interval. Cell lysates were sonicated for 5 min and centrifuged at 14,000 rpm at 4 °C for 15 min. Protein concentrations were determined by BCA Protein Assay Kit (Pierce™, Cat #23225). Then, lysates were mixed with RPPA lysis buffer and 4X SDS/B-Me sample buffer (40% Glycerol, 8% SDS, 0.25 M Tris-HCL, pH 6.8 and Beta-mercaptoethanol (B-Me) added before use at 1/10) to have final concentration of 1.22 μg/μl for further process by MD Anderson Center.

Cell lysate samples were serially diluted two-fold for 5 dilutions (undiluted, 1:2, 1:4, 1:8; 1:16) and arrayed on nitrocellulose-coated slides in an 11×11 format to produce sample spots. Sample spots were then probed with antibodies by a tyramide-based signal amplification approach and visualized by DAB colorimetric reaction to produce stained slides. Stained slides were scanned on a Huron TissueScope scanner to produce 16-bit tiff images. Sample spots in tiff images were identified and their densities quantified by Array-Pro Analyzer.

For data processing, relative protein levels for each sample were determined by interpolating each dilution curve produced from the densities of the 5-dilution sample spots using a “standard curve” (SuperCurve) for each slide (antibody). SuperCurve is constructed by a script in R, written by Bioinformatics. Relative protein levels are designated as log2 values (RawLog2). All relative protein level data points were normalized for protein loading and transformed to linear values, which are designated “Normalized Linear” (NormLinear). “Normalized Linear” values were transformed to log2 values (NormLog2) and then median-centered for hierarchical clustering analysis (NormLog2_MedianCentered). Comparison between different treatment groups and the corresponding control group (vehicle-treated empty vector (EV, GDNF-) group in MCF-7 or vehicle-treated non-targeting control (shNTC, GDNF-) group in SUM44 models, respectively) was accomplished by limma package in R, with results sorted by adjusted p value (Table S2). Top 40 significant (adjusted p value < 0.05) targets were selected and plotted into heatmap ranked by LogFC in RET-OE + GDNF group (shNTC + GDNF group in SUM44). Heatmaps of all significant targets in hierarchical clustering analysis could be found in Figure S6.

### Western Blotting (WB)

Cells were seeded in 6-well plates and were subjected to desired treatment at 70-80% confluency. Cells were all pre-incubated in low serum medium (0.5% FBS medium for MCF-7, T-47D and MM134 and 0.5% CSS medium for SUM44) for 16 hr (serum starvation) followed by 30 min of GDNF treatment. Pralsetinib (Blueprint Medicine, GAVRETO™) was provided not only in the low serum medium during serum starvation, but also applied during GDNF (or distilled water) treatment, DMSO as the vehicle control. Specially, 10 ng/ml GFRα1 (R&D Systems, Cat #714-GR-100) was applied together with GDNF for T-47D.

After rinsing the cells with cold PBS, cell lysates were harvested in 1.5X RIPA buffer (50 mM Tris, pH 7.4, 150 mM NaCl, 1 mM EDTA, 0.5% Nonidet P-40 (IGEPAL), 0.5% Sodium Deoxycholate, 0.1% SDS) with 1X Halt Protease and Phosphatase Inhibitor Single-Use Cocktail (100X, Thermo, Cat # 78442) added. The lysates were sonicated for 5 min and centrifuged at 14,000 rpm at 4 °C for 15 min. Protein concentrations were determined by BCA Protein Assay Kit (Pierce™, Cat #23225). 50 µg protein was loated to each well of a 15-well 8% denaturing gel, and was transferred in Tris-Glycerol buffer with 20% Methanol at 90 V for 90 min.

Both Li-Cor Odyssey Western Blotting protocol and Bio-Rad General Western Blotting protocol were followed as adaptive to Odyssey® Imaging System (Li-Cor) and ChemiDoc XRS+ Imaging System (Bio-Rad), respectively. Briefly, as described in Li-Cor Odyssey Western Blotting protocol, PVDF membrane was blocked in PBS blocking buffer (Fisher Scientific, Li-Cor, Cat # NC1660556) for 1 hr, and incubated in 1:1000 diluted primary antibody at 4 °C overnight. Membrane was washed in 0.1% PBS-T and incubated with conjugated secondary antibody (1:10000, IRDye® 800CW Goat anti-Rabbit IgG (H + L), Li-Cor, Cat #926-32211 or IRDye® 680LT Goat anti-Mouse IgG (H + L), Li-Cor, Cat #925-68020) for 55 min at RT. After washing, the membrane was imaged by Li-Cor Odyssey® Imaging System.

As for a Bio-Rad General Western Blotting protocol, PVDF membrane was blocked in 5% BSA diluted in the washing buffer, 0.1% TBS-T, instead. In this method, HRP-conjugated secondary antibodies were used (see Table 2.3). Membrane was finally developed using Clarity™ Western ECL Substrate (Bio-Rad, Cat #1705061) and imaged by ChemiDoc XRS+ Imaging System (Bio-Rad) or exposed to films which were developed through Konica SRX-101A Film Processor. Of note, phospho-RET (pRET) bands were not able to be visualized following the Odyssey WB protocol so all the pRET blots were generated from the second method. To strip and re-probe the blots, 1X NewBlot Stripping buffer (Li-Cor, Cat #928-40032) was used standardly following the manufacturer’s protocol.

## Notes

### Competing Interest Statement

The authors have declared no competing interest.

