## Supplementary Materials for "Targeting RET in Brain Metastases from Estrogen Receptor Positive Breast Cancer"

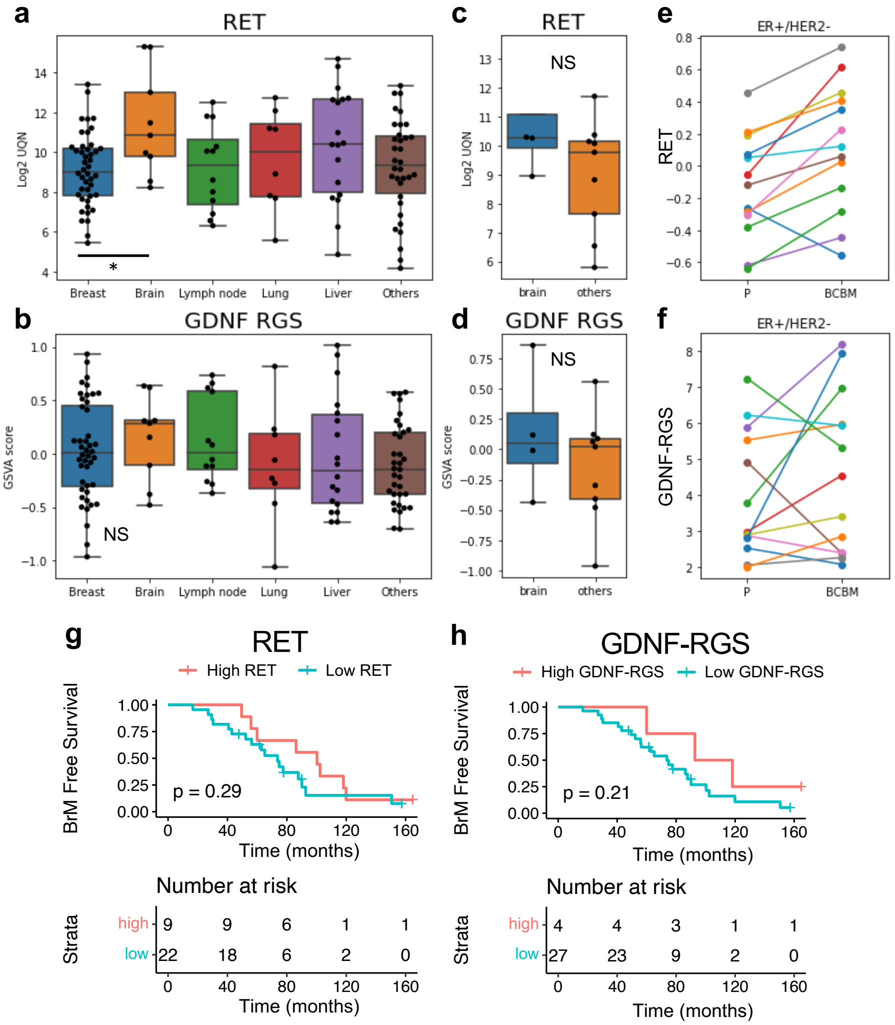


**Figure S1. RET expression and signaling was increased in breast cancer brain metastasis.**

(a-b) RET expression (log2NormCPM:log2 normalized counts per million) (a) and GSVA score of GDNF-RGS signature (b) in primary (n = 40) and metastatic breast cancer samples (n = 70) from AURORA dataset. Wilcoxon test. (c-d) RET expression (c) and GSVA score (d) of primary tumors grouped by future metastatic sites from ER+ subgroup of AURORA dataset. (e-f) Paired ladder plot of RET expression (e) and GDNF/RGS GSVA score (f) in BrM versus matched primary tumor. (g) KM plot of brain metastasis free survival with high/low GDNF-RGS GSVA scores in ER+ cases of METABRIC. (h) KM plot of brain metastasis free survival with high/low RET expression in ER+ cases of METABRIC.

**Figure S2. Pralsetinib potently inhibits RET activation and RET-mediated cell proliferation in breast cancer cell line models.**

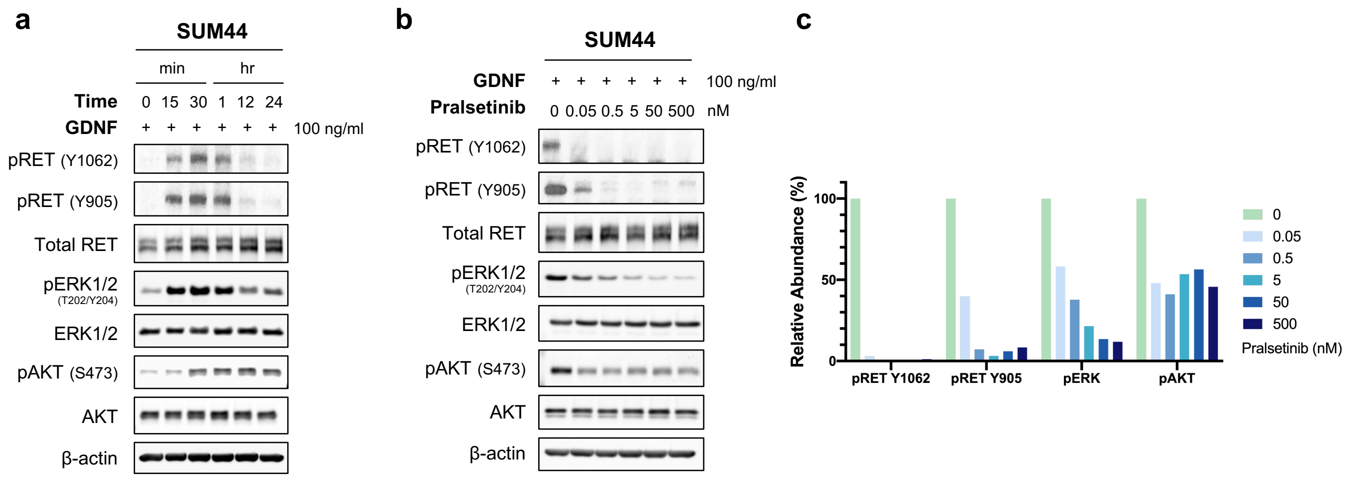


**d**

**ULA Dose Response**

| Cell lines | GDNF | IC50 (μM) |
|---|---|---|
| MCF-7 EV | - | 3.66 |
| MCF-7 EV | + | $3.12 \times 10^{-3}$ |
| MCF-7 RET-OE | - | 5.74 |
| MCF-7 RET-OE | + | $4.34 \times 10^{-2}$ |

**Figure S2. RET expression and signaling in ER+ breast cancer cell line models.**
(a) GDNF time course response in SUM44 cell model. (b-c) Immunoblotting (b) and quantification (c) of pralsetinib dose response in SUM44 cells. (d) IC50 calculated in ULA dose response assay of Pralsetinib (refer to Figure 3c). Cells were treated with 100 ng/ml GDNF for 30 min in (b-c). 50 ug protein is loaded per well. shNTC; cells transfected with non-targeting control shRNA

**Figure S3. RET knockdown reduces enhancement of cell proliferation and sphere formation mediated by GDNF.**

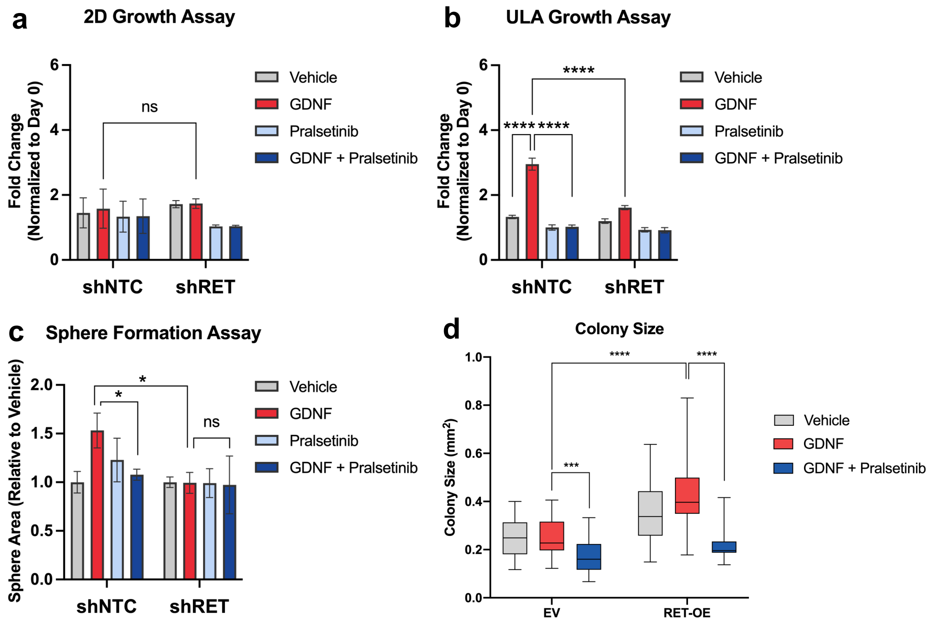


**Figure S3. RET knockdown reduces enhancement of cell proliferation and sphere formation mediated by GDNF.**

(a-b) Cell growth assay in 2D (a) and ULA (b) conditions in MCF-7 RET knockdown models. (c) Quantification of sphere formation assay. (d) Quantification of colony size in soft agar assay. GDNF, 100 ng/ml. Pralsetinib, 500 nM. Vehicle, distilled water for GDNF, and DMSO for Pralsetinib. EV; empty vector. RET OE; RET overexpressing cells. shNTC; cells transfected with non-targeting control shRNA. shRET; cells transfected with shRNA targeting hRET. *, $0.01 < p < 0.05$, **, $0.001 < p < 0.01$, ***, $0.0001 < p < 0.001$, ****, $p < 0.0001$.

**Figure S4. Development of brain metastases through intracardiac injection of MCF-7 in NSG mice.**

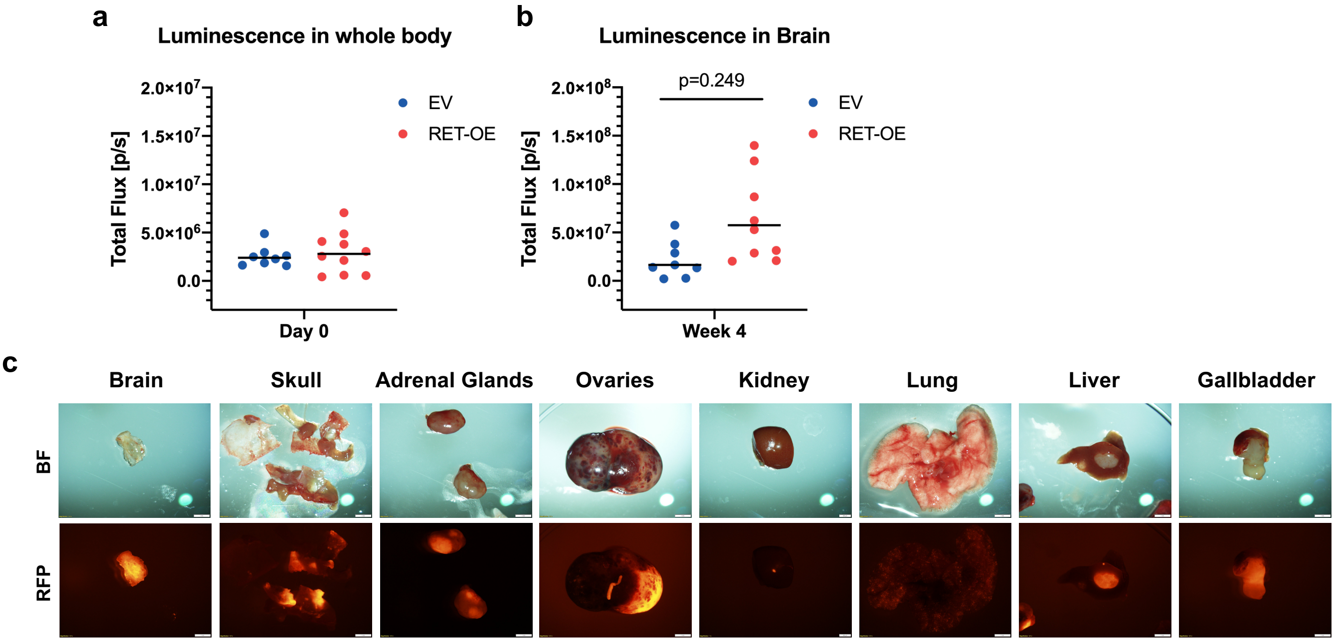


**Figure S4. Development of brain metastases through intracardiac injection in NSG mice.**

(a) Quantification of luminescence in the whole animal body right after injection. (b) Quantification of luminescence in the brain at week 4. Luminescence was measured 5 min after i.p. injection of D-luciferin (150 mg/kg). All luminescence was measured 5 min after i.p. injection of D-luciferin (150 mg/kg). (c) Representative bright field (BF) and RFP 1X dissecting scope images of brain mets and macromets in other sites. EV; animals injected with RFP-labeled empty vector transfected MCF-7 cells. RET-OE; animals injected with RFP-labeled RET overexpressing MCF-7 cells.

**Figure S5. Development of brain metastases through orthotopic injection of SUM44PE in NSG mice.**

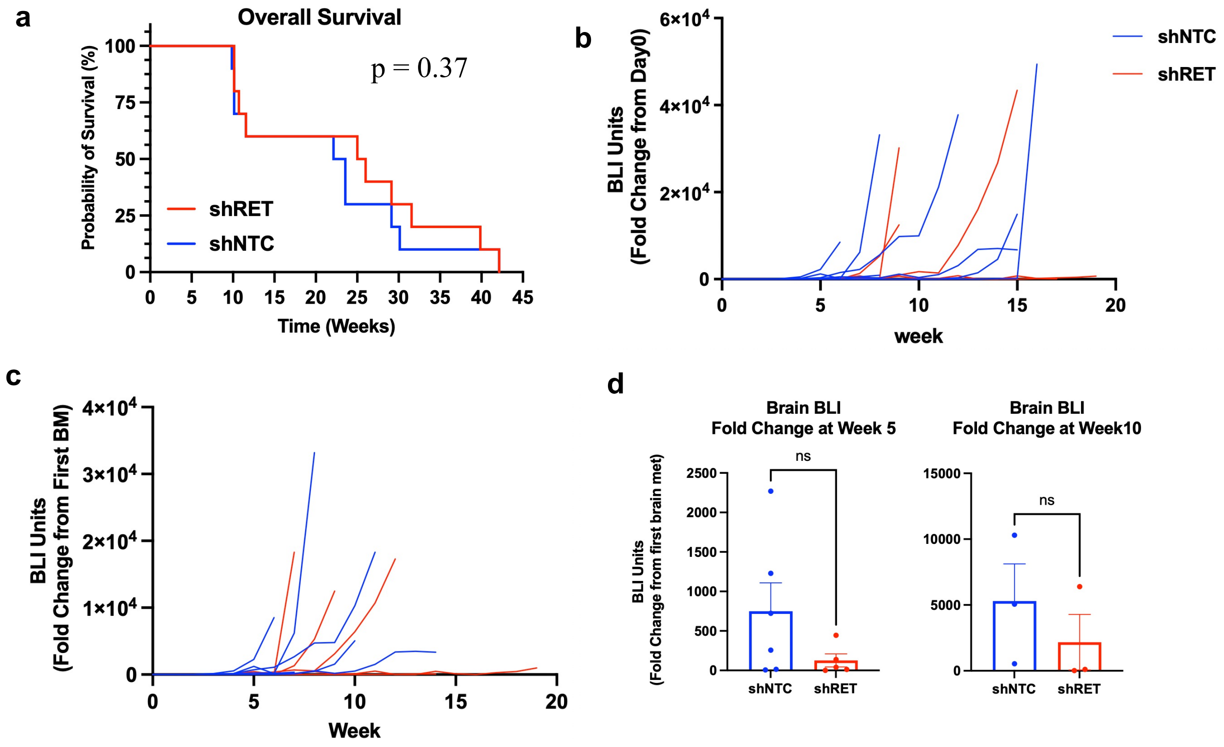


(a) Overall survival. Competing risk analysis (Gray's test). (b) Quantification of luminescence in individual brains throughout the whole experiment. Normalized to the first day when the primary tumor was removed. (c) Quantification of luminescence in the brain for who develops brain metastasis. Normalized to the first brain metastasis signal. (d) Quantification of luminescence in the brain at early (week 5) and late (week 10) stage of progression using IVIS imaging. Normalized to the first brain metastasis signal.

**Table S1. Survival rate of orthotopic injection of SUM44PE in NSG mice.**

| Group | # of BrM | Rate (%) |
|---|---|---|
| shNTC | 6(7) | 85.7% |
| shRET | 5(6) | 83.3% |

**Figure S6. Hierarchical clustering of significant targets identified in Human Phospho-Kinase Array and RPPA.**

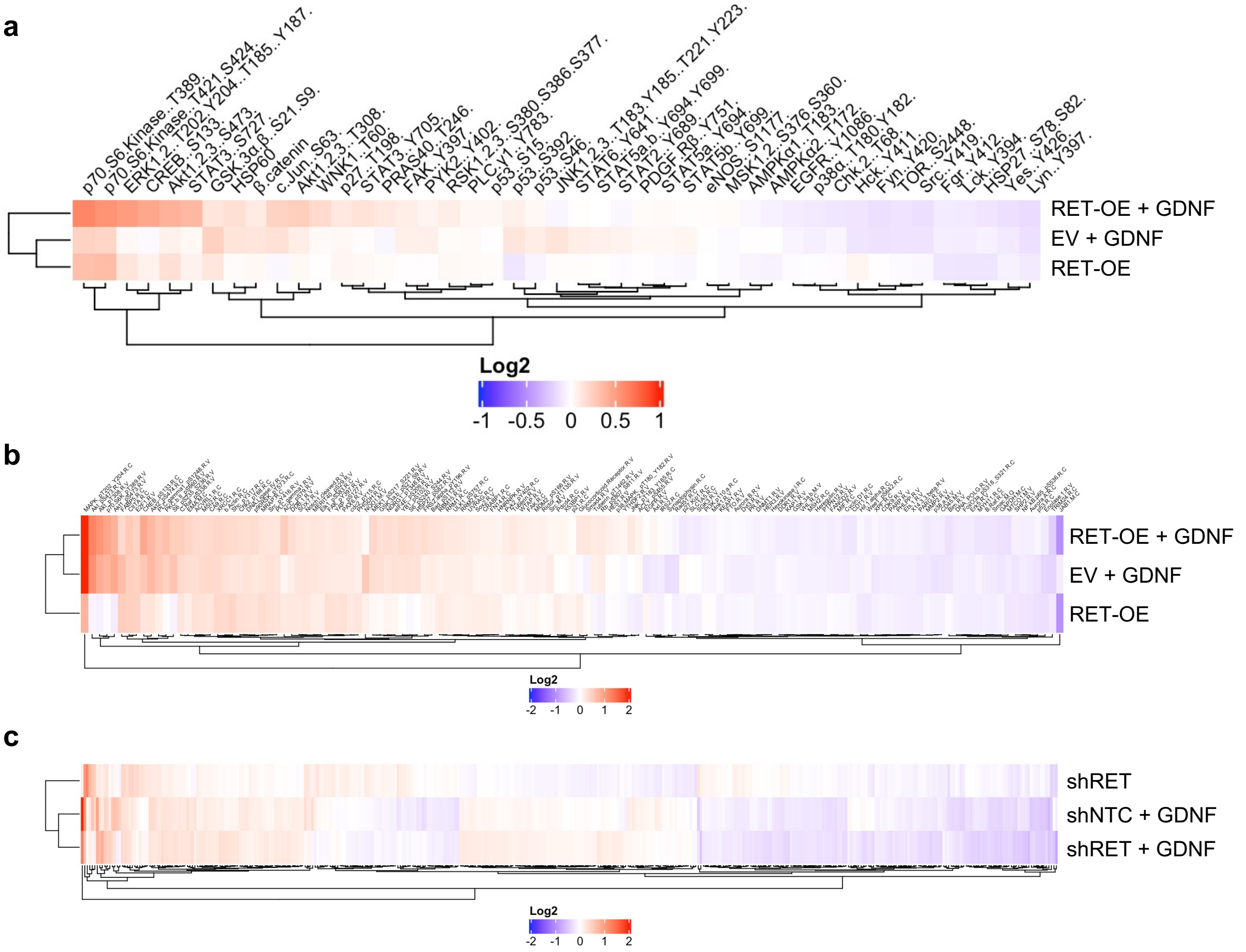


**Figure S5. Hierarchical clustering of significant targets identified in Human Phospho-Kinase Array and RPPA.**
(a) Hierarchical clustering of significant targets in RET-overexpressing MCF-7 cell models identified in Human Phospho-Kinase Array. (b-c) Hierarchical clustering of significant targets identified in RPPA using MCF-7 (b) or SUM44 (c) cell models. Cells were treated with 100 ng/ml GDNF for 30 min. EV; empty vector transfected cells. RET-OE; RET overexpressing cells. shNTC; cells transfected with non-targeting control shRNA. shRET; cells transfected with shRNA targeting shRET.

**Figure S7. PLCγ-1 knockdown does not attenuate GDNF-mediated cell migration.**

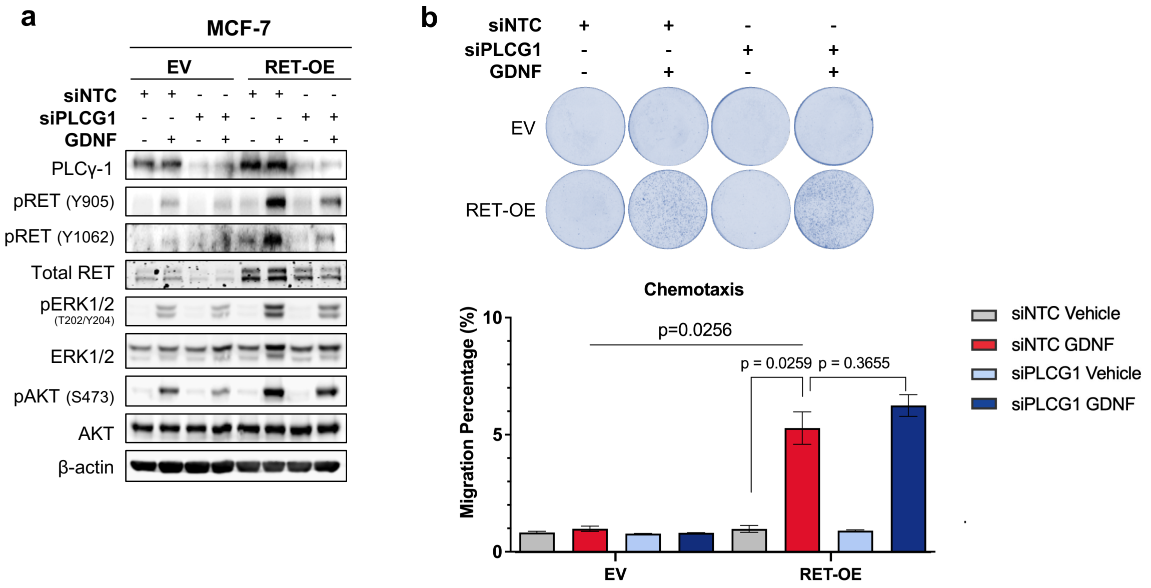


(a). Immunoblotting in empty vector or RET overexpressing MCF-7 cell lines. ERK1/2 and AKT are GDNF-responsive RET downstream targets that can be downregulated by PLCγ-1 knockdown. (b) Images and quantification of crystal violet–stained transwell inserts showing the migration of RET overexpressing MCF-7 cells transfected with siPLCG1 or non-targeting control RNA in response to GDNF chemotaxis. One way ANOVA was performed to test for statistical significance. GDNF, 100 ng/ml, and vehicle is distilled water for GDNF. EV; empty vector. RET-OE; RET overexpressing cells. NTC; non targeting control RNA.
